# *Tropomyosin 1* genetically constrains *in vitro* hematopoiesis

**DOI:** 10.1101/631895

**Authors:** CS Thom, CD Jobaliya, K Lorenz, JA Maguire, A Gagne, P Gadue, DL French, BF Voight

## Abstract

Identifying causal variants and genes from human genetics studies of hematopoietic traits are important to enumerate basic regulatory mechanisms underlying these traits, and could ultimately augment translational efforts to generate platelets and/or red blood cells *in vitro*. To identify putative causal genes from these data, we performed computational modelling using available genome-wide association data sets for platelet traits. Our model identified a joint collection of genomic features enriched at established platelet trait associations and plausible candidate variants. Additional studies associating variation at these loci with change in gene expression highlighted the *Tropomyosin 1* (*TPM1*) among our top-ranked candidate genes. CRISPR/Cas9-mediated *TPM1* knockout in human induced pluripotent stem cells (iPSCs) enhanced hematopoietic progenitor development, increasing total megakaryocyte as well as erythroid cell yields. Our findings may help explain human genetics associations and identify a novel genetic strategy to enhance *in vitro* hematopoiesis.

## Introduction

Elucidating genetic mechanisms governing hematopoiesis has broad value in understanding blood production and hematologic diseases (Ulirsch et al., 2019). Given interest in generating platelets and red blood cells (RBCs) from *in vitro* culture of induced pluripotent stem cells (iPSCs) (An et al., 2018; Giani et al., 2016; Ito et al., 2018) there is also translational value in harnessing genetic and molecular processes that regulate hematopoiesis. Cost-effective blood cell generation will require novel strategies based on better knowledge of underlying mechanisms driving *in vitro* development.

*In vitro* hematopoietic systems might be improved by identifying novel factors from human genetic studies. Genome wide association studies (GWAS) have linked hundreds of single nucleotide polymorphisms (SNPs) with platelet and/or red cell trait variability (Astle et al., 2016; Gieger et al., 2011). Because most GWAS SNPs are non-coding, likely influencing transcriptional expression of key genes (Hindorff et al., 2009; Tak and Farnham, 2015), it has been challenging to derive functional biochemical understanding of the key genes of action (Edwards et al., 2013; Tak and Farnham, 2015; Xu and Taylor, 2009). Relatively few studies have elucidated biochemical mechanisms for blood trait variability loci (Nurnberg et al., 2012; Pleines et al., 2017; Polfus et al., 2016; Simon et al., 2016; Soranzo et al., 2009). One strategy to narrow focus on candidate genes is to link non-coding variation to expression of nearby genes (Ritchie et al., 2014; Shihab et al., 2015; Ulirsch et al., 2019). However, for platelet trait variation alone, GWAS have thus far implicated >6700 expression quantitative trait loci (eQTL) affecting expression of >1100 genes (Methods). Hence, there is a clear need to more specifically identify putatively functional sites.

Actin cytoskeletal dynamics play key roles in hematopoiesis (Lambert, 2015; Standing, 2017; Thon et al., 2010). Tropomyosin proteins coat most actin filaments and regulate actin functions (Gunning and Hardeman, 2017; Meiring et al., 2018). All four human tropomyosin genes (1-4) are expressed in human hematopoietic cells, and *Tropomyosin 4* promotes platelet development (Pleines et al., 2017). Genetic studies have also suggested a role for *Tropomyosin 1* (*TPM1*) in human platelet trait variation (Astle et al., 2016), though no prior studies have elucidated if or how *TPM1* impacts human hematopoiesis.

Here, we utilized penalized regression to construct a model that predicted platelet GWAS associations based on epigenetic data sets as features for the prediction. Our model built from platelet trait GWAS loci reliably distinguished sentinel GWAS SNPs, as well as platelet-relevant genes and loci. Among these prioritized sites were SNPs that regulate *TPM1* expression. To validate this putative candidate gene and to explore its function, we used CRISPR/Cas9 genome editing to discover that cultured *TPM1*- deficient induced pluripotent stem cells enhanced hematopoietic progenitor cell formation. In turn, this increased functional megakaryocyte yield. Thus, our framework stratified relevant functional loci, and helped identify *TPM1* manipulation as a novel strategy to enhance *in vitro* hematopoiesis.

## Results

### Penalized regression model identifies genetic regulatory loci for hematopoiesis

GWAS have linked hundreds of single nucleotide polymorphisms (SNPs) with variability in human platelet traits (Astle et al., 2016). To focus our studies on credible functional follow-up candidates, we utilized a penalized logistic regression framework, i.e., the least absolute shrinkage and selection operator (LASSO) (Tibshirani, 1996; Zou and Hastie, 2005), using 860 features to construct a model that distinguished platelet trait GWAS SNPs from control SNPs matched for allele frequency, distance to gene, and number of SNP proxies in strong linkage disequilibrium (Figure 1A, Methods, and Supplementary File 1).

**Figure 1.**
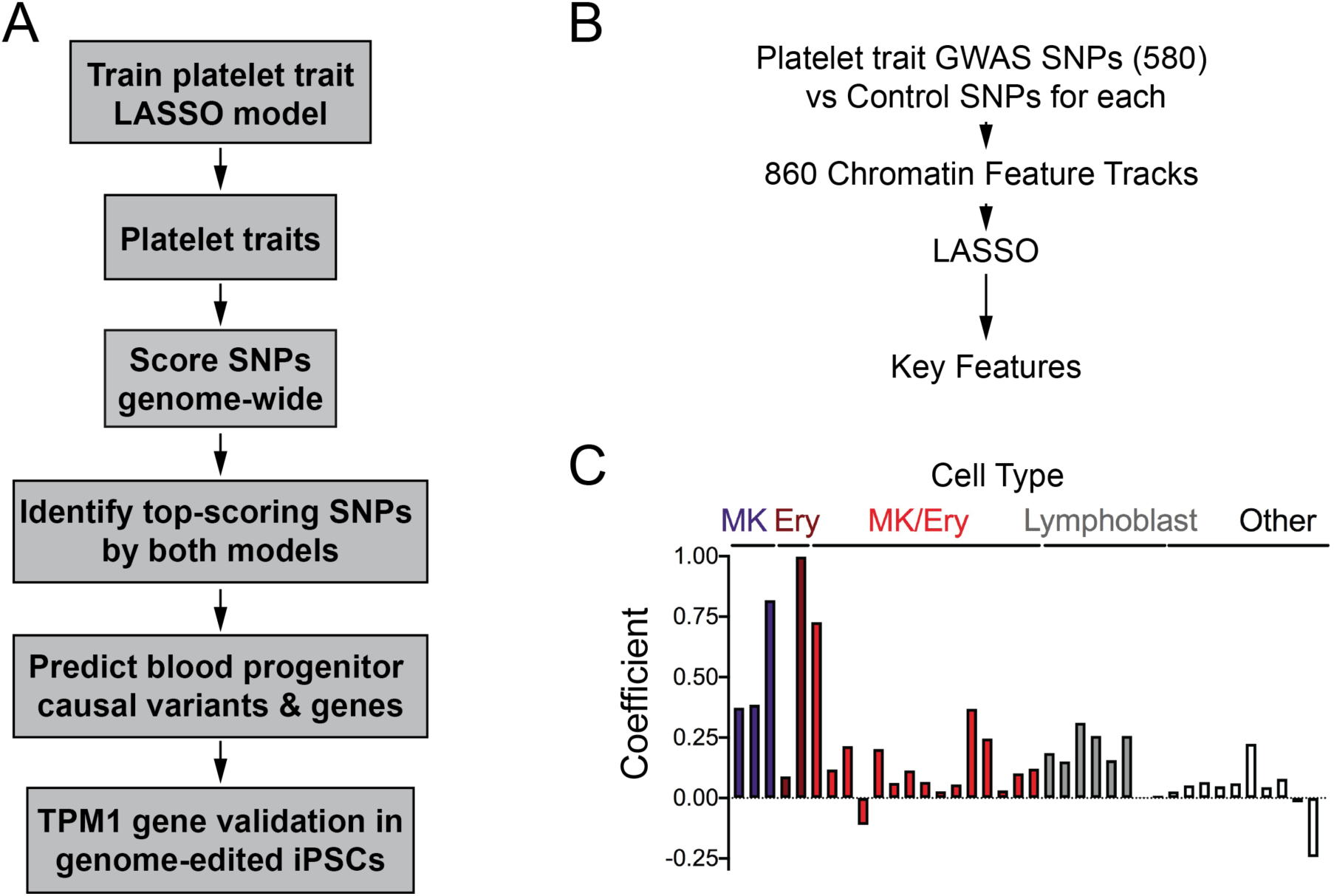
A penalized regression-based approach to identify hematopoietic regulatory loci and genes. **(A)** Schematic outline of our approach. We generated a penalized regression-based predictive scoring algorithm based on platelet trait GWAS loci, and applied the resultant scoring algorithm genome-wide to predict causal variants and genes. We validated this model computationally and through validation of *TPM1* function in induced pluripotent stem cells (iPSCs). **(B)** To generate a penalized regression model, 580 platelet trait GWAS SNPs (Astle et al., 2016) and matched control SNPs (∼100 per GWAS SNP (Schmidt et al., 2015)) were analyzed for overlap with 860 chromatin features (e.g., histone marks and TF binding sites). **(C)** Penalized regression (LASSO) (Tibshirani, 1996) analysis identified 38 chromatin features from the indicated cell types that best discriminated GWAS SNPs. Bar heights are LASSO coefficients, indicating the relative importance of each feature. MK, primary megakaryocytes. Ery, peripheral blood derived erythroblasts. MK/Ery, K562 cells. Lymphoblast, GM12878 or GM12891.

Our ‘platelet trait model’ was trained on 580 genome-wide-significant platelet trait-related SNPs from a large recent GWAS of human blood trait variation (Astle et al., 2016), along with 860 chromatin features (Figure 1B). These GWAS SNPs affected human platelet count (PLT), platelet-crit (PCT), mean platelet volume (MPV), and/or platelet distribution width (PDW). For each GWAS SNP, we identified matched control SNPs based on distance to nearest gene, number of SNPs in linkage disequilibrium, and minor allele frequency. Model performance in the training phase was assessed using standard approaches (i.e., 10-fold cross-validation).

The resultant predictive model comprised 38 epigenomic features that best distinguished platelet trait GWAS SNPs from controls (Figure 1C, Figure 1-figure supplement 1 and Supplementary File 2). This model discriminated positive and negative labeled examples with an Area Under the Receiving Operator Curve (AUC) = 0.726 (Figure 1-figure supplement 1).

### Genome-wide model application

We calculated trait-enrichment scores genome-wide based on SNP overlap with each of the selected features, weighted by our penalized regression model coefficients (Methods and Supplementary File 2). As expected, SNP scores were significantly higher for platelet trait GWAS SNPs relative to SNPs genome-wide (p<0.0001 by ANOVA, Figure 2A). While some care in interpretation was required, it was encouraging that the model selected biologically plausible features. GATA1, GATA2, SCL and FLI1 are critical hematopoietic transcription factors (Pimkin et al., 2014; Tijssen et al., 2011), and most of our features came from hematopoietic cell types (primary MK, peripheral blood-derived erythroblasts, K562 with MK/erythroid potential, and GM12878/GM12891 lymphoblasts; Supplementary File 2).

**Figure 2.**
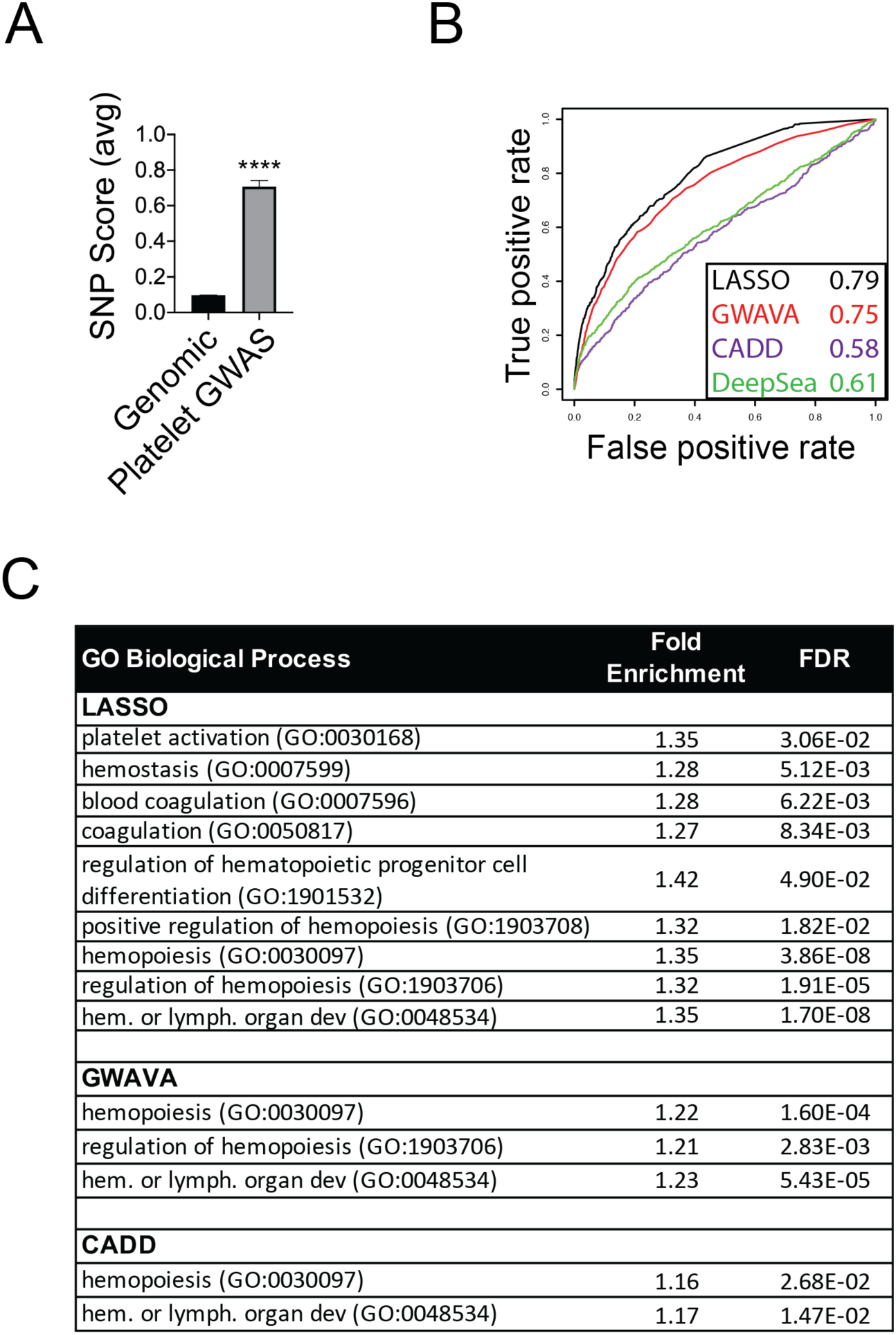
Penalized regression model identifies genes relevant to platelet and hematopoietic biology. **(A)** SNP scores for platelet model training SNPs were significantly higher than genome-wide SNP scores. Bars represent mean+-SEM, ****p<0.0001. **(B)** Performance comparison of our platelet trait model to DeepSEA (Zhou and Troyanskaya, 2015), GWAVA (Ritchie et al., 2014), and CADD (Kircher et al., 2014) for platelet trait SNP identification. AUC values are shown in the legend. **(C)** Platelet and hematopoiesis pathways (Ashburner et al., 2000) identified by the highest-scoring (top 1%) SNPs genome-wide for the indicated models, excluding established platelet trait loci (Astle et al., 2016) (FDR, False Discovery Rate).

### Application of additional prediction methods

Our goal was to use a compilation of methods and evidence to specify loci with high functional likelihood. Some models have been previously developed to identify active genomic loci (e.g., CADD (Kircher et al., 2014), GWAVA (Ritchie et al., 2014), and DeepSEA (Zhou and Troyanskaya, 2015)). We compared the effectiveness of these models, and our trait-specific model, to discriminate platelet trait GWAS sites from a set of ∼10,000 randomly selected SNPs. We used AUC values to assess model performance. Our trait-specific model had the highest AUC in this analysis (0.79), and GWAVA also performed well (AUC 0.75, Figure 2B).

GWAVA prioritizes functional impact of non-coding genomic elements without regard for lineage or trait specificity (Ritchie et al., 2014). Hence, our results suggested that chromatin marks associated with active gene regulatory regions were enriched in platelet trait GWAS loci.

However, hematopoiesis- and blood lineage-specific chromatin regulatory mechanisms are also critical for blood development (Aranda-Orgilles et al., 2016; Heuston et al., 2018; Petersen et al., 2017a). It was difficult to parse hematopoietic biological rationale in the regulatory elements prioritized by GWAVA scoring. Therefore, we pursued further validation of our trait-specific model, in an effort to best specify loci and related genes that were important for hematopoiesis, megakaryopoiesis, and/or platelet biology.

### Genome-wide model validation

Encouraged by the features we selected and our model performance, we next sought to derive external support for the model selected by our regression framework. First, we evaluated the biological specificity of variation prioritized by the model. This was particularly important, given practical limitations associated with fine-mapping and cellular validation experiments. Gene Ontology analysis of the top 1% highest-scoring SNPs indicated that the nearest genes to penalized regression-prioritized variants were enriched for biologically relevant pathways, even after removing GWAS-significant sites (Figure 2C and Supplementary Files 3-5). While many pathways related to platelet function and coagulation were associated, generalized hematopoiesis- and hematopoietic progenitor cell-related pathways were also included.

Second, we asked whether our SNP scores correlated with summary association statistics for platelet trait-GWAS data (Astle et al., 2016). Indeed, variants that were nominally associated with platelet traits but did not reach genome-wide significance and not included in our model (P-value between 0.05 and 5×10^-8^) had significantly higher average scores compared to SNPs that were not obviously associated (P-value > 0.05, Figure 2-figure supplement 1). This correlation suggested that our scoring algorithm was valid genome-wide and could potentially reveal true biological associations, as had the GWAS itself (Gieger et al., 2011; Nurnberg et al., 2012; Simon et al., 2016; Soranzo et al., 2009).

Finally, we asked if regulatory gene enhancer regions were enriched with high scoring SNPs by our model, consistent with regulatory function. We found that our model assigned higher scores to SNPs in FANTOM5 enhancer regions (Andersson et al., 2014) compared with other sites genome-wide, consistent with the hypothesis that functional non-coding SNPs associate with active regulatory regions (Farh et al., 2014; Tak and Farnham, 2015) (Figure 2-figure supplement 2, enhancer region scores >0.9 vs genome-wide baseline <0.1). We further observed that enhancer regions in hematopoietic cell types scored significantly higher than enhancers from irrelevant control cells (Figure 2-figure supplement 2). These data suggest trait specificity in hematopoietic enhancers, consistent with prior studies (Petersen et al., 2017b), and the broader hypothesis about tissue-specific trait heritability as reported elsewhere (Finucane et al., 2015; Trynka et al., 2015). Collectively, our findings indicated that we could successfully target hematopoietic and platelet trait-relevant loci.

### Exemplary candidate locus and gene identification

Next, we used computational predictions, including our own model, to stratify sites and related genes for functional validation. Given practical limitations related to follow-up validation, we wanted to narrow our focus to a modest number of loci (e.g., <20). We reasoned that functional SNPs would (i) be in high linkage disequilibrium (LD) with established platelet trait GWAS loci, (ii) score highly relative to other SNPs within that LD block, (iii) regulate target gene(s) as expression quantitative trait loci (eQTL), and (iv) overlap GATA binding sites (Grant et al., 2011; Mathelier et al., 2016). We prioritized GATA binding sites based on the importance of GATA factors in hematopoiesis (Freson et al., 2011; Tijssen et al., 2011) and in our penalized regression model (Supplementary File 2). We specifically focused our attention on sites scored in the top 5% genome-wide by our platelet trait model and a more generalized machine learning-based model that performed also well in validation analyses (GWAVA (Ritchie et al., 2014), Figure 2B).

This stratification approach identified 15 loci and related genes, including SNPs known to impact hematopoiesis, megakaryocyte and/or platelet biology (Table 1 and Figure 2-figure supplement 3). In principle, *any* site meeting these stringent criteria could form the basis for interesting biological follow up experiments.

**Table 1.** Penalized regression-based fine-mapping identifies eQTLs in established platelet trait GWAS loci that overlie GATA binding sites. Listed SNPs are within platelet trait GWAS LD blocks (EUR r^2^>0.7), scored in the top 5% by our platelet trait model and by GWAVA (Ritchie et al., 2014), overlap canonical or near-canonical GATA binding sites, and are eQTLs for at least 1 gene (GTEx V7) (Ardlie et al., 2015). Associated gChromVAR scores are shown (Ulirsch et al., 2019). Genes in **bold** have known hematopoietic function. SNP rsIDs and locations refer to hg19 genome.

| rsID (hg19) | Chr | Pos (Mb) | Platelet score (%ile) | GWAVA score (%ile) | gChromVAR (PP PLT) | Nearest Gene | eQTL Gene(s) |
| --- | --- | --- | --- | --- | --- | --- | --- |
| rs11240368 | 1 | 205.1 | 1.12 (97th) | 0.52 (95th) |  | *DSTYK* | *CNTN2*, *TMEM81* |
| rs3771535 | 2 | 70.0 | 0.94 (95th) | 0.53 (95th) | 0.01 | *ANXA4* | *GMCL1*, *SNRNP27* |
| rs10180681 | 2 | 121.0 | 1.41 (98th) | 0.63 (97th) | 0.01 | *RALB* | *EPB41L5*, *PTPN4*(Gu and Majerus, 1996), ***RALB***(Clough et al., 2002) |
| rs10180682 | 2 | 121.0 | 1.41 (98th) | 0.64 (97th) | 0.01 | *RALB* | *EPB41L5*, *PTPN4*(Gu and Majerus, 1996), ***RALB***(Clough et al., 2002) |
| rs9646785 | 2 | 172.0 | 1.27 (98th) | 0.58 (96th) |  | *TLK1* | *GAD1*, *GORASP2* |
| rs6771578 | 3 | 167.4 | 1.14 (97th) | 0.6 (96th) | 0.003 | *PDCD10* | ***PDCD10***(Cohen et al., 2019), *SERPINI1*, *WDR49* |
| rs12652692 | 5 | 77.8 | 3.62 (99th) | 0.57 (96th) | 0.01 | *LHFPL2* | *LHFPL2*, *SCAMP1* |
| rs72793280 | 5 | 131.6 | 2.73 (99th) | 0.89 (99th) | 0.001 | *P4HA2* | *ACSL6*, ***P4HA2***(Gilkes et al., 2013; Wang et al., 2019), ***PDLIM4***(Bauer et al., 2000), *SLC22A4*, *SLC22A5* |
| rs1741820 | 6 | 122.8 | 1.75 (99th) | 0.55 (96th) |  | *HSF2* | *HSF2*, *PKIB* |
| rs342293 | 7 | 106.4 | 1.80 (99th) | 0.94 (99th) | 0.99 | *CC71L* | ***PIK3CG***(Soranzo et al., 2009)* |
| rs13265995 | 8 | 56.7 | 1.75 (99th) | 0.6 (96th) |  | *TMEM68* | ***LYN***(Ming et al., 2011; Suzuki-Inoue et al., 2002), *TGS*, *TMEM68* |
| rs9704108 | 11 | 0.3 | 1.12 (97th) | 0.85 (99th) | 0.087 | *IFITM2* | *IFITM2* |
| rs11071720 | 15 | 63.3 | 1.53 (98th) | 0.58 (96th) | 0.98 | *TPM1* | *APH1B*, *LACTB*, *RAB8B*, ***TPM1***(Gieger et al., 2011)** |
| rs2316513 | 17 | 2.0 | 0.92 (95th) | 0.54 (95th) | 0.005 | *EST1A* | *DPH1*, *SMG6*, ***SRR***(Yakovenko et al., 2018) |
| rs1654439 | 19 | 55.6 | 2.69 (99th) | 0.58 (96th) | 0.002 | *RDH13* | ***GP6***(Quach et al., 2018; Suzuki-Inoue et al., 2002), *NLRP2*, ***RDH13***(Eicher et al., 2016) |

\*eQTL in human platelets (Soranzo et al., 2009), but not in GTEx tissues (Ardlie et al., 2015).
\*\*Function suggested by *D. rerio* morpholino experiments (Gieger et al., 2011).

Two of these loci stood out as high-scoring variants by the recently described gChromVAR algorithm (Ulirsch et al., 2019), which is based on accessible chromatin regions in hematopoietic cells (Table 1). First, rs342293 is a GWAS SNP (Gieger et al., 2011) that regulates *PIK3CG* gene expression (Soranzo et al., 2009) and lies within accessible chromatin in hematopoietic progenitor cell types (Corces et al., 2016a) (Figure 3A-B). The GATA site is disrupted in the presence of the SNP minor allele (Figure 3C). In platelets, *PIK3CG* activity regulates PIK3 signaling (Hawkins et al., 2014) and response to collagen (Pasquet et al., 2000). Individuals harboring this minor allele had increased MPV and decreased platelet reactivity (Soranzo et al., 2009) (Figure 3D).

**Figure 3.**
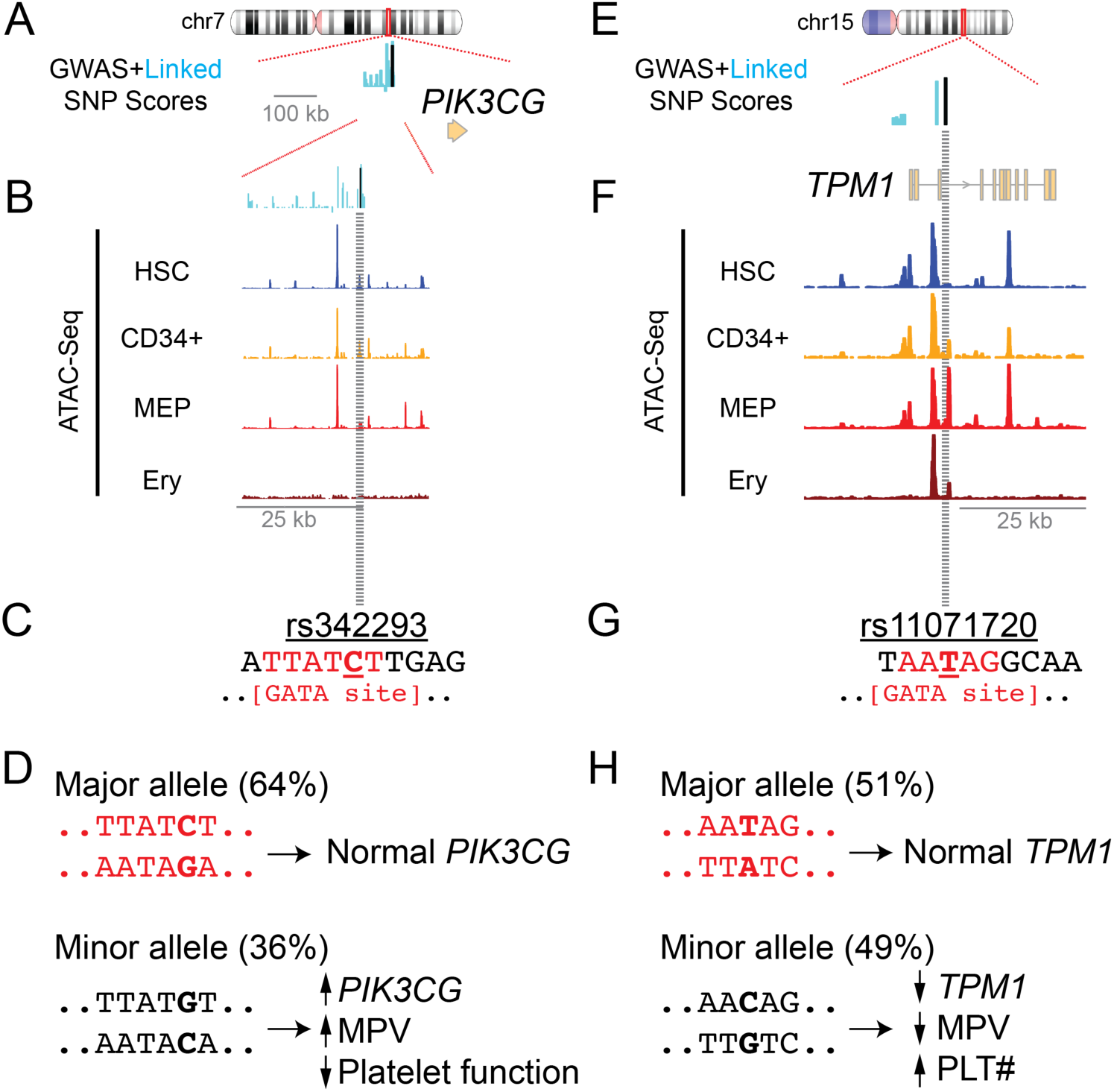
Exemplary high-scoring eQTLs near the *PIK3CG* and *TPM1* gene loci overlap putative GATA binding sites and are associated with altered platelet traits. **(A)** High-scoring platelet trait GWAS SNP rs342293 (Gieger et al., 2011) (black) and linked SNPs (EUR r^2^>0.7, cyan) lie upstream of *PIK3CG*. Bar heights depict SNP scores. **(B)** This region overlaps a dynamic accessible chromatin region during hematopoiesis (Corces et al., 2016b). ATAC-Seq data are shown for hematopoietic stem cells (HSC), CD34+ hematopoietic progenitor cells, megakaryocyte-erythroid progenitors (MEP), and erythroblasts (Ery). **(C)** The local DNA sequence for rs342293 (underlined) includes a canonical GATA binding site (Mathelier et al., 2016) (red). **(D)** Platelet phenotypes associated with rs342293 alleles (Soranzo et al., 2009). **(E)** SNP scores near platelet trait GWAS SNP rs11071720 (Astle et al., 2016) (black) and linked SNPs (EUR r^2^>0.7, cyan). Bar heights depict SNP scores. **(F)** Accessible chromatin (ATAC-Seq) regions at this locus are shown for hematopoietic stem cells (HSC), CD34+ hematopoietic progenitor cells, megakaryocyte-erythroid progenitors (MEP), and erythroblasts (Ery) (Corces et al., 2016b). **(G)** Local DNA sequence shows a putative GATA binding site (Mathelier et al., 2016) (red) around rs11071720 (underlined ‘T’). **(H)** The major and minor rs11071720 alleles and associated platelet phenotypes (Astle et al., 2016; Fehrmann et al., 2011). Allele percentages based on UCSC genome browser and dbSNP.

A second variant, rs11071720, found within the 3^rd^ intron of the *Tropomyosin 1* (*TPM1*) gene locus, also attracted our attention. This sentinel GWAS SNP scored highly compared to linked SNPs (EUR r^2^>0.7) and overlapped accessible chromatin in hematopoietic cells (Corces et al., 2016b) (Figure 3E-F). The rs11071720 minor allele, which disrupts a near-canonical GATA binding site, is an eQTL associated with decreased *TPM1* expression (Ardlie et al., 2015; Fehrmann et al., 2011), higher platelet count, and lower MPV (Astle et al., 2016) (Figure 3G-H and Figure 3-figure supplement 1).

Tropomyosin proteins regulate actin cytoskeletal functions, which are critical for hematopoietic, megakaryocyte, and platelet biology (Italiano et al., 2007; Lambert, 2015; Pleines et al., 2017; Standing, 2017). Although morpholino studies showed *TPM1* to be important for zebrafish thrombopoiesis (Gieger et al., 2011), no prior study had examined the effect of *TPM1* during human hematopoiesis. Based on these and the human genomics data, we hypothesized that *TPM1* is an important effector of hematopoiesis and ultimately platelet biology. Thus, in what follows, we focus our cellular validation studies on *TPM1*, under the hypothesis that this SNP regulated the expression of this gene.

### Tropomyosin 1 modulation enhances in vitro hematopoiesis

We investigated functions for the *TPM1* gene in an *in vitro* human model of primitive hematopoiesis (Sim et al., 2017). We expected that total gene deletion would show stronger effects than non-coding SNP modification (Sankaran and Orkin, 2013). Using CRISPR/Cas9, we targeted a ∼5kb region containing *TPM1* exons 4-8 in iPSCs (Figure 4A), anticipating creation of a null allele (Schevzov et al., 2011). We confirmed deletion by sequencing and western blot (Figure 4B-C and Figure 4-figure supplement 1). In total, we obtained 3 *TPM1* knockout (KO) clones from 2 separate genetic backgrounds. Karyotype and copy number variation analyses confirmed that engineering these clones did not introduce any *de novo* genomic aberrancies (data not shown).

**Figure 4.**
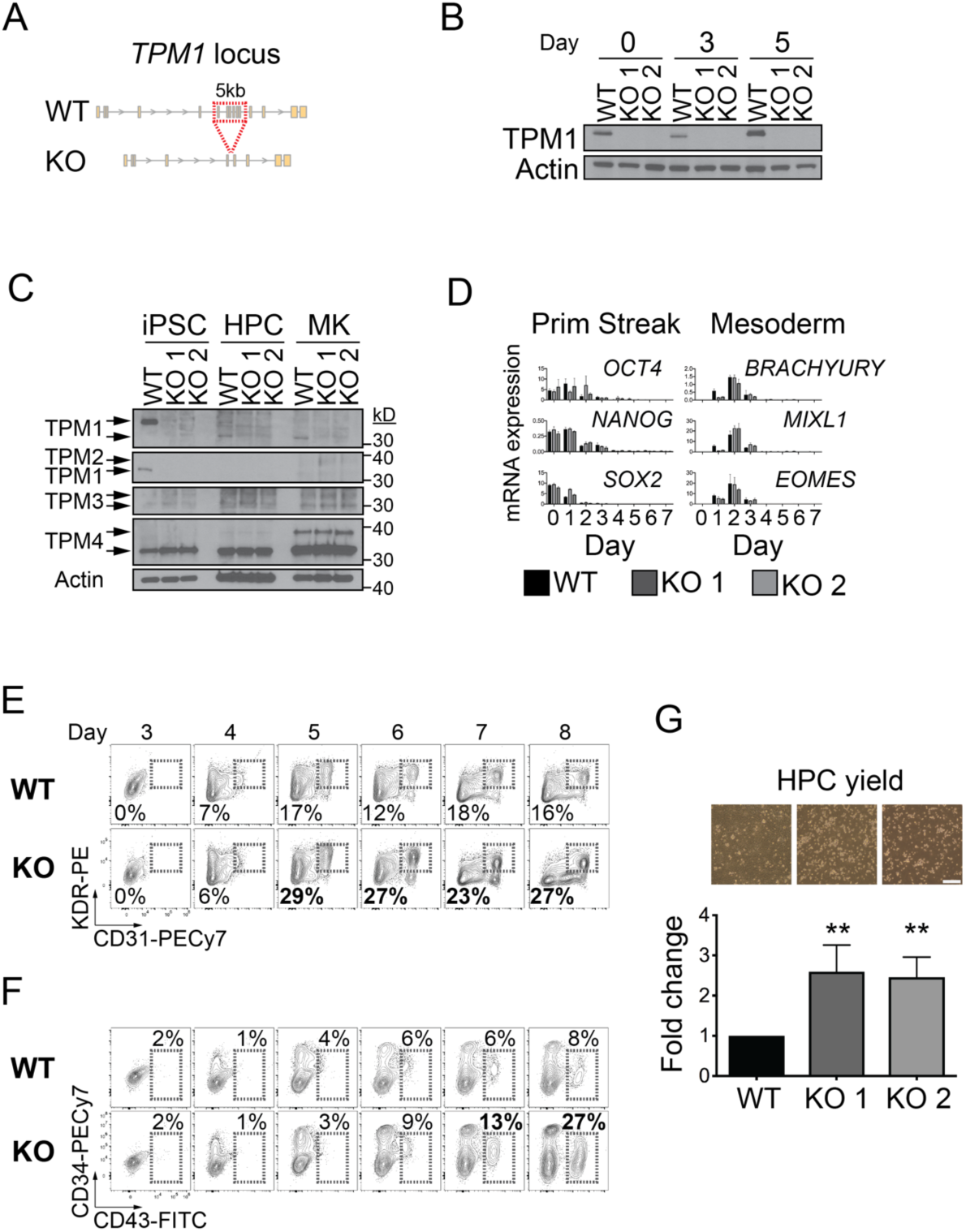
*TPM1* deficiency enhances iPSC-derived hematopoietic progenitor cell formation. **(A)** A 5 kb region (*TPM1* exons 4-8, red box) was targeted for CRISPR/Cas9-mediated deletion to create KO iPSCs. **(B)** Western blots showing TPM1 protein expression at the indicated differentiation days. **(C)** Western blots showing TPM1-4 expression in wild type (WT) and KO iPSCs, hematopoietic progenitor cells (HPC, differentiation d8) and FACS-sorted MKs (CD41^+^/CD42b^+^, expansion d3). TPM1 antibodies targeted exon 4 (top) or exon 9d (2^nd^ panel). **(D)** Primitive streak and mesoderm gene expression are normal in differentiating KO iPSCs. **(E,F)** KO cells yield (**E**) KDR^+^/CD31^+^ cells and (**F**) CD43^+^ hematopoietic progenitor cells (HPC) with normal kinetics, but in enhanced abundance. Percent (%) cells within boxed regions for WT and KO clone 2 are shown from a representative experiment. **(G)** Quantification of WT and KO non-adherent HPCs on differentiation day 8. Bars represent fold change in HPCs (mean±SD) vs WT for ≥4 experiments. (Top) Culture images on differentiation d8, with HPCs (light color) floating above an adherent monolayer. Scale bar, 20mm.

TPM1 protein was present during early iPSC differentiation, but downregulated non-adherent hematopoietic progenitor cells and differentiated MKs (Figure 4B-C). Early differentiation proceeded normally in KO clones, with normal patterns of primitive streak and mesoderm gene expression (Figure 4D), as well as pluripotency marker loss (Figure 4-figure supplement 1). The kinetics by which KDR^+^/CD31^+^ endothelial/hemogenic endothelial cells and CD43^+^ hematopoietic progenitor cells (HPCs) emerged were also normal (Figure 4E-F). In this culture system, KDR^+^/CD31^+^ cells include both HPC precursor cells (hemogenic endothelium) as well as cells destined for a purely endothelial fate.

Unexpectedly, we found that KO cultures enhanced generation of KDR^+^/CD31^+^ as well as CD43^+^ HPCs (Figure 4E-F). We quantified HPC abundance by cell counting and flow cytometry, observing that KO HPC yield doubled that of WT controls (Figure 4G). We confirmed this finding in a KO clone from a genetically distinct iPSC background (Figure 4-figure supplement 2). All HPCs retained normal hematopoietic cell surface marker expression (Figure 4-figure supplement 3).

Next, we investigated whether KO HPCs would yield functional megakaryocytes in increased quantities. Liquid expansion culture revealed normal mature CD41^+^/CD42b^+^ megakaryocyte yield per HPC (Figure 5A). With twice as many starting HPCs, this meant that total megakaryocyte recovery increased ∼2-fold in KO cultures. KO megakaryocyte morphology was normal (Figure 5-figure supplement 1), and megakaryocyte activation in response to agonists were normal-to-increased (Figure 5B). Microarray gene expression analyses of WT and KO megakaryocytes revealed no statistically significant changes in megakaryocyte genes (Figure 5-figure supplement 2 and Supplementary File 6).

**Figure 5.**
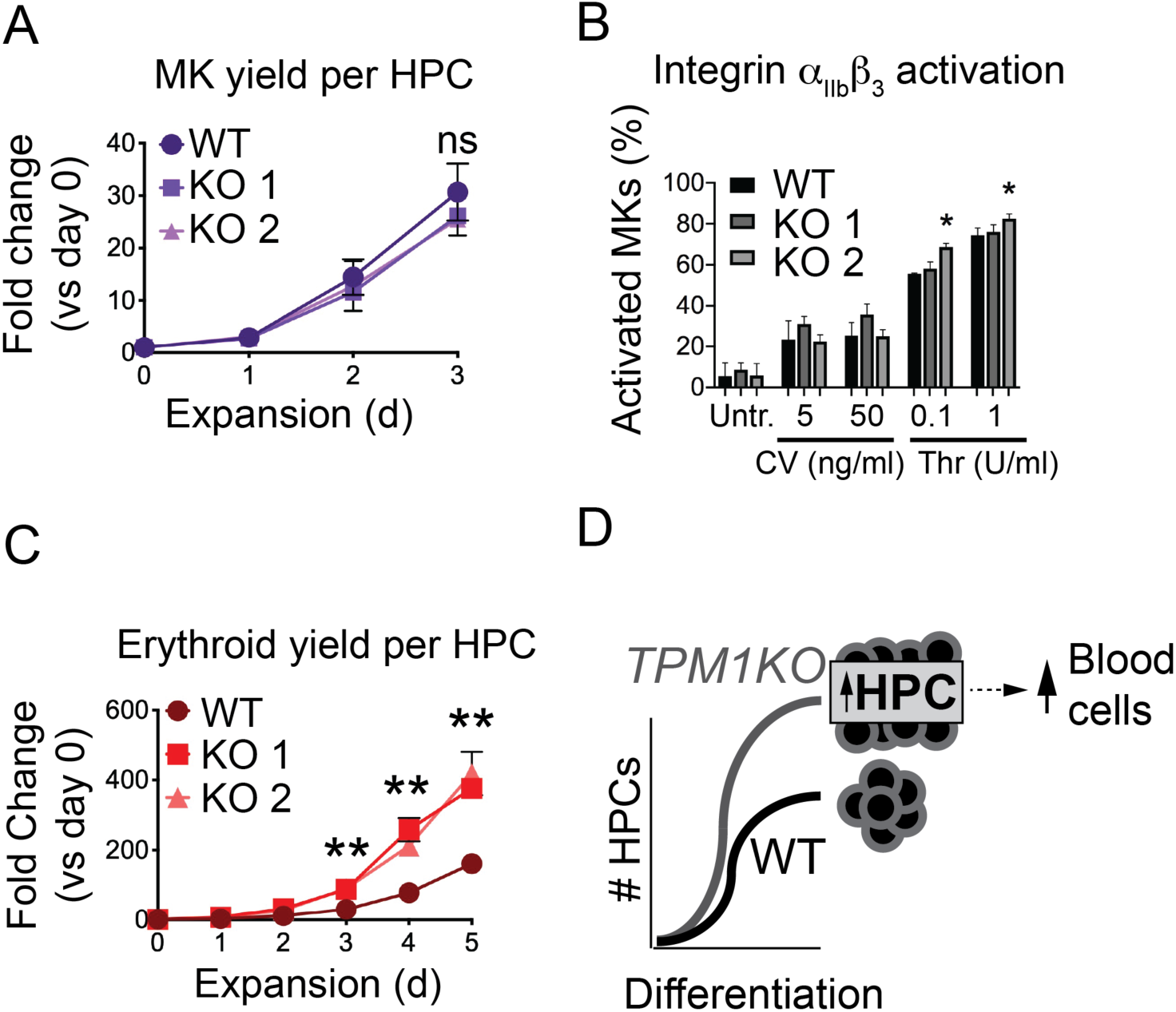
TPM1-deficient hematopoietic progenitor cells yield normal-to-increased quantities of functional, mature hematopoietic cells. **(A)** WT and KO HPCs put into MK expansion culture generate equivalent numbers of MKs. Points represent CD41^+^/CD42b^+^ MKs percentage multiplied by total cell count, normalized to cell count on day 0. **(B)** *TPM1* KO MKs respond appropriately to platelet agonists. WT and KO MKs were incubated in the presence of Convulxin (CV) or Thrombin (Thr) at the indicated concentrations, and the percentage of activated MKs (PAC-1^+^/CD41^+^/CD42b^+^) were quantified. *p<0.05 by ANOVA vs WT. **(C)** KO HPCs put into erythroid expansion culture generate more erythroid cells than WT HPCs. Points represent CD235^+^ percentage multiplied by total cell count, normalized to cell count on day 0. **(D)** Model in which KO iPSCs yield more HPCs than WT, generating more total MKs and RBCs. ns, not significant, **p<0.01 by ANOVA.

The early hematopoietic phenotype in KO cultures was unexpected. We asked whether KO HPCs might also enhance yield of other blood cell types. Indeed, KO HPCs spawned normal-to-increased quantities of erythroid and myeloid cells (Figure 5C and Figure 5-figure supplement 3). Hence, *TPM1* deletion enhanced formation of HPCs with multilineage potential (Figure 5D).

### Tropomyosin 1 locus is prioritized by red cell trait-based penalized regression model

We were surprised by the early hematopoietic effects of *TPM1* deletion, given that rs11071720 has only been genetically linked with platelet traits (Astle et al., 2016). We therefore investigated whether this finding could have been predicted using human genetics data. We found that rs11071720-linked regulatory variants were marginally associated with red cell traits, although these data did not meet genome-wide significance (Figure 5-figure supplement 4). It is possible that future studies with improved power will reveal a true statistical association with red cell traits at this locus.

We also trained an additional model for red cell traits, using an analogous framework and regulatory features as described for platelet traits (Methods), but instead using 818 red blood cell trait-related GWAS SNPs affecting red blood cell count (RBC count), hematocrit (HCT), mean red cell corpuscular volume (MCV), and/or red cell distribution width (RDW). The resultant model included 78 features and distinguished red cell trait GWAS SNPs with an AUC = 0.732 (Figure 5-figure supplement 5 and Supplementary File 7). When used as a scoring algorithm genome-wide, this red cell trait model displayed performance similar to the platelet trait model (Figure 5-figure supplement 6, Figure 5-figure supplement 7 and Supplementary File 8).

Interestingly, our red cell model scored rs11071720 in the 96^th^ percentile genome-wide (Supplementary File 9). This prioritization agrees with *TPM1* impacting both megakaryocyte and erythroid lineages. The other 14 sites that scored in the top 5% by both platelet and red cell models might also be expected to regulate early hematopoietic biology, and could form the basis for future cellular validation experiments (Supplementary File 9). Indeed, several of these genes are known to regulate hematopoiesis.

## Discussion

Genetic insights could augment efforts to generate blood products *in vitro* (An et al., 2018; Giani et al., 2016; Ito et al., 2018), but relatively few genetically implicated loci or genes have been functionally validated (Nurnberg et al., 2012; Pleines et al., 2017; Polfus et al., 2016; Simon et al., 2016; Soranzo et al., 2009). The purposes of our present study were to establish (i) whether computational approaches using available genomic data could prioritize trait-specific sites and genes that impact hematopoiesis, megakaryopoiesis and/or platelet biology, and (ii) to validate the function of a novel candidate gene (i.e. *TPM1*) in a translationally-relevant iPSC model. Our data support a model whereby *TPM1* deficiency enhances *in vitro* formation of multilineage HPCs (Figure 5D). In addition to understanding a genetic modifier of hematopoietic traits (Astle et al., 2016), application of our results may augment *in vitro* megakaryocyte and erythroid cell yields.

Broadly, the successful implementation of this trait-specific penalized regression method demonstrates a tunable approach to variant and gene identification. Our pipeline is similar to prior methods that have stratified loci based on chromatin feature data (e.g. GWAVA (Ritchie et al., 2014) and fGWAS (Pickrell, 2014)), but is readily scalable to any set of loci and chromatin features. For blood-related traits, it is an adaptable complement to established and excellent scoring models such as gchromVAR (Ulirsch et al., 2019).

Given the scope of the present study, the most important functional result was enhanced yield of HPCs and functional megakaryocytes. Our results were directionally consistent with human genetic data (Astle et al., 2016), finding that decreased *TPM1* expression portends higher megakaryocyte yield. The molecular mechanism(s) driving enhanced HPC formation will be of considerable biologic and translational interest, and such studies are ongoing. *TPM1* KO-related increases in HPC formation may complement or synergize previously described approaches that enhanced later stages of hematopoiesis (Giani et al., 2016; Ito et al., 2018; Thon et al., 2014; Wen et al., 2012).

Early hematopoietic function for *TPM1* was unexpected based on blood genetics (Astle et al., 2016). Our model may have prioritized some ‘early’ hematopoietic sites, given that many chromatin features derived from relatively immature megakaryocytes (Tijssen et al., 2011) as well as K562 cells, which can act as progenitors for erythroid or megakaryocyte lineages. Indeed, some of the sites targeted general hematopoietic- and HPC-related pathways (Figure 2C). Chromatin feature data from mature megakaryocytes may enable future models to more specifically target late stage megakaryopoiesis and/or platelet sites. Alternatively, *TPM1* could have separate function early and late in hematopoiesis, akin to *GATA2* (Castaño et al., 2019).

Though a lack of robust detection methods precluded accurate platelet production quantitation in our culture system, normal function of derived megakaryocytes suggests an overall increase in megakaryocytes yield would translate into higher platelet production. Importantly, our findings do not exclude additional effects on terminal megakaryopoiesis or erythroid development *in vitro*, nor *in vivo* effects outside the scope of our iPSC model.

Enhanced hematopoiesis in *TPM1KO* iPSCs contrasts detrimental effects of *TPM1* deficiency on organism fitness in other contexts (Anyanful et al., 2001; Gieger et al., 2011; Rethinasamy et al., 1998). For example, abrogated *D. rerio* thrombopoiesis with *tpma*-directed morpholinos (Gieger et al., 2011) resembles human *TPM4* deficiency (Pleines et al., 2017) rather than *TPM1* deficiency. This highlights the importance of species-specific genetic validation, particularly given inter-species disparities in hematopoiesis (Pishesha et al., 2014).

In conclusion, using a penalized regression modeling approach to functional variant identification led us to define a role for *TPM1* in constraining *in vitro* hematopoiesis. Recent advances increasing per-MK platelet yields (Ito et al., 2018) have focused a spotlight on increasing cost effectiveness of *in vitro* MK generation. In addition to improved recognition of genes and mechanisms underlying quantitative hematopoietic trait variation, application of the computational approach described herein could also help to specify trait-specific causal genetic variants for virtually any clinically relevant human trait.

## Methods

### In silico analyses

Relevant data sets and coding scripts can be found on GitHub (https://github.com/thomchr/2019.PLT.TPM1.Paper). Human genome version hg19 was used for all analyses, and we utilized the LiftOver script when necessary (https://bioconductor.org/packages/release/workflows/html/liftOver.html). GWAS summary statistics are publicly available (http://www.bloodcellgenetics.org/).

### Expression Quantitative Trait Locus analysis

To estimate the number of eQTLs implicated by prior platelet trait GWAS, SNPs in high LD with established GWAS loci(Astle et al., 2016) (EUR r^2^>0.9) were identified using PLINK. From this set of SNPs, eQTLs and affected genes were identified from GTEx V7 (Ardlie et al., 2015). Numbers reported in the text reflect unique eQTL SNPs, which often functioned across multiple tissues. The affected gene estimate reflects the number of unique Ensembl gene identifiers (ENSG).

### SNP selection

From a total of 710 genome-wide significant GWAS SNPs (p<5E-8) affecting platelet count, platelet-crit, mean platelet volume, and/or platelet distribution width (Astle et al., 2016), 580 comprised our platelet model training SNP set. These 580 had rsIDs that were recognized by the <u>G</u>enomic <u>R</u>egulatory <u>E</u>lements and <u>G</u>WAS <u>O</u>verlap algo<u>R</u>ithm (GREGOR) (Schmidt et al., 2015) tool, which we used to select control SNPs based on Distance to nearest gene, number of SNP LD proxies linked to the lead associated SNP (r^2^ ≥ 0.8), and minor allele frequency. We identified ∼2000 matched controls for each training SNP, all with a minor allele frequency >10%. This minor allele cutoff was necessary to limit the effects of very low control SNP frequencies on the resultant model.

From a total of 1003 genome-wide significant GWAS SNPs (p<5E-8) affecting red cell count, hematocrit, mean corpuscular volume, and/or red cell distribution width (Astle et al., 2016), 818 had rsIDs recognized by GREGOR. These comprised the red cell model training SNP set. We identified ∼2000 matched controls with minor allele frequency >10% for each training SNP.

### Chromatin feature selection

We collected a subset of available features tracks from ENCODE (Feingold et al., 2004), including data for hematopoietic (K562, GM12878 and GM12891) as well as other cell types (e.g., H1-hESC, HUVEC, HeLa, HepG2, etc.). We also collected available feature tracks from primary MKs and hematopoietic cells (Paul et al., 2013; Tijssen et al., 2011). The only modification to any of these genomic data sets was peak-calling in MK-derived chromatin immunoprecipitation-sequencing (ChIP-Seq) tracks (Zhang et al., 2008). See Supplementary File 1 for a list of these features.

### Penalized regression modeling

To generate our model, we first analyzed training set GWAS SNPs and matched controls SNPs for overlap with 860 chromatin features (data set available on GitHub). Columns representing our 3 baseline parameters (Distance to Nearest Gene, number of LD proxies linked to the lead associated SNP, and Minor Allele Frequency) were also included in this data table for each SNP. This chromatin feature overlap data file was then analyzed using the least absolute shrinkage and selection operator (LASSO, L1 regularization, glmnet version 2.0-18) (Tibshirani, 1996; Zou and Hastie, 2005) with 10-fold cross-validation and forced inclusion of the 3 baseline parameters. Baseline parameters were assigned penalty factors of 0 (to force inclusion), while other chromatin features were assigned penalty factors of 1. Features and coefficients were taken from the λ_se_. There were 41 features included in our platelet model and 81 features in our red cell model, including the 3 baseline parameters. For downstream genome-wide analyses, we scored all SNPs within NCBI dbSNP Build 147 based on coefficients and overlaps with model features.

### Model performance comparison

We used public databases to obtain SNP scores for alternative models (CADD v1.3 (Kircher et al., 2014), GWAVA unmatched score (Ritchie et al., 2014), DeepSEA (Zhou and Troyanskaya, 2015); https://cadd.gs.washington.edu/download, http://www.sanger.ac.uk/resources/software/gwava, http://deepsea.princeton.edu). For each model, we identified scores for platelet trait GWAS SNPs and a random selection of 10,000 control SNPs. We then used ROCR (Sing et al., 2005) to compare model performance in discriminating GWAS SNPs from controls, and report the area under the receiver operating characteristic (AUC) for each model. An analogous pipeline was used to analyze the ability of each model to discriminate red cell trait-related GWAS SNPs from controls.

For specific sites, including rs11071720, we obtained gChromVAR scores (Ulirsch et al., 2019) (https://molpath.shinyapps.io/ShinyHeme/).

### Model Evaluation

To assess biological specificity, we identified the top 1% highest-scoring SNPs from each model (platelet model, red cell model, GWAVA, CADD) after excluding all red cell or platelet trait-associated GWAS loci. We then used closestBed (https://bedtools.readthedocs.io/en/latest/content/tools/closest.html) to identify the nearest gene to each of these SNPs. Genes and positioned were defined by BioMart (http://www.biomart.org/). We then used the Gene Ontology resource (http://geneontology.org/) to analyze pathway enrichment. Input analysis settings were Binomial tests and calculated FDR for GO Biological Process complete. Pathways identified with FDR<5% are presented in Figure 2E and Supplementary Files 4-7.

Enhancer regulatory regions were defined according to the FANTOM5 data set (Andersson et al., 2014). Presented FANTOM5 data represent scores for all overlapping SNPs from dbSNP 147.

### Linkage disequilibrium structure assessment

The SNP Annotation and Proxy Search tool (https://archive.broadinstitute.org/mpg/snap/ldsearch.php), LDlink (https://analysistools.nci.nih.gov/LDlink), and 1000 Genomes Project (phase 3) data were used to measure linkage disequilibrium in the EUR population.

### Transcription factor binding site identification

To identify GATA sites, the genomic sequence context for SNPs of high interest were obtained using the UCSC Table Browser (Kent et al., 2002) and analyzed for matches by manual curation of canonical or near-canonical GATA binding motif in all orientations (AGATAA, TTATCA, AATAGA, TTATCT; GATAA, AATAG, CTATT, TTATC).

### Human iPSC generation

iPSC models were generated as described from peripheral blood mononuclear cells (Maguire et al., 2016). The “CHOP10” and “CHOP14” lines were used in this study. CRISPR/Cas9-mediated genome editing was performed as described (Maguire et al., 2019) per protocols from the CHOP Human Pluripotent Stem Cell Core Facility (https://ccmt.research.chop.edu/cores_hpsc.php) with the following guide sequences: 5’ (1) ATGACGAAAGGTACCACGTCAGG, 5’ (2) TGAGTACTGATGAAACTATCAGG, 3’ (1) CCCTTTTCTTGCTGCTGTGTTGG, 3’ (2) GGAGAGTGATCAAGAAATGGAGG.

Karyotyping (Cell Line Genetics, Madison, WI) and copy number variation (CHOP Center for Applied Genomics, Philadelphia, PA) analyses were performed per institutional protocols.

### iPSC hematopoietic differentiation and analysis

iPS cell cultures and primitive hematopoietic differentiations were performed as per published protocols (Mills et al., 2014, 2013; Paluru et al., 2013; Sim et al., 2017). iPS cells were maintained on irradiated mouse embryonic feeder cells in human embryonic stem cell (ESC) medium (DMEM/F12 with 20% knockout serum, 100 μM non-essential amino acids, 0.075% sodium bicarbonate, 1 mM sodium pyruvate, 2 mM glutamine, 50 U/ml penicillin, 50 g/ml streptomycin (all from Invitrogen), 10−4 M β–mercaptoethanol (Sigma, St Louis, MO), and 10 ng/ml human bFGF (Stemgent)). Medium was changed at least every 2 days and colony clusters passaged weekly to new feeders ESC medium containing ROCK inhibitor (10 μM) using TrypLE (Invitrogen) and gentle scraping.

About 1 week prior to differentiation, iPSCs were transitioned to a ‘feeder-free’ state by culturing on Matrigel-coated wells (BD Biosciences; 6-well tissue culture plate, Falcon 3046) in ESC medium under atmospheric O_2_ conditions.

Throughout hematopoietic differentiation, cells were maintained at 37°C in 5% CO_2_, 5% O_2_ and 90% N_2_. All media were supplemented with 2 mM glutamine, 50 μg/ml ascorbic acid (Sigma, St. Louis, MO), 150 μg/ml transferrin (Roche Diagnostics), and 4 × 10^−4^ M monothioglycerol (Sigma). Media and cytokines were changed daily as follows (Gadue et al., 2006): Days 0-1 RPMI (Invitrogen) with 5 ng/ml BMP4, 50 ng/ml VEGF and 25 ng/ml Wnt3a; Day 2 RPMI with 5 ng/ml BMP4, 50 ng/ml VEGF and 20 ng/ml bFGF; Day 3 SP34 (Invitrogen) with 5 ng/ml BMP4, 50 ng/ml VEGF and 20 ng/ml bFGF; Days 4-5 SP34 with 15 ng/ml VEGF and 5 ng/ml bFGF; Day 6 serum free differentiation medium (SFD) with 50 ng/ml VEGF, 100 ng/ml bFGF, 100 ng/ml SCF, and 25 ng/ml Flt3L; Days 7-9 SFD with 50 ng/ml VEGF, 100 ng/ml bFGF, 100 ng/ml SCF, 25 ng/ml Flt3L, 50 ng/ml TPO, 10 ng/ml IL-6, and 0.05-2 U EPO. In all differentiations, marked cell death occurred through day 2, after which time surviving cells formed an adherent monolayer. Analyses during differentiation therefore used 0.25% trypsin-EDTA (ThermoFisher Scientific; 1 ml/well, 5 mins at room temperature) to dissociate monolayer cells.

By day 6-7, non-adherent floating hematopoietic progenitor cells (HPCs) appeared. HPCs were collected on days 7-9 and either frozen or used directly for further culture and/or analyses. HPCs cultured in 50 ng/ml thrombopoietin and 25 ng/ml SCF to generate megakaryocytes, 2U erythropoietin and 25 ng/ml SCF to generate erythroid cells, or 200 ng/ml granulocyte/macrophage colony stimulating factor to generate myeloid cells.

Flow cytometry gating strategies for pluripotency (SSEA3^+^/SSEA4^+^), hemogenic endothelium (KDR^+^/CD31^+^), hematopoietic progenitors (CD43^+^ and CD41^+^/CD235^+^) and terminal lineages have been previously validated (Mills et al., 2014, 2013; Paluru et al., 2013; Sim et al., 2017).

### Flow cytometry

Flow cytometry analysis was performed on a Cytoflex LX and FACS-sorting was performed on a FACS Aria II (BD Biosciences). Flow cytometry data were analyzed using FlowJo 10 (Tree Star, Inc.). The following antibodies were used for flow cytometry: FITC-conjugated anti-CD41 (BioLegend), PE-conjugated anti-CD42b (BD Biosciences), APC-conjugated anti-CD235 (BD Biosciences), PB450-conjugated anti-CD45 (BioLegend), AF488-conjugated anti-SSEA3 (BioLegend, AF647-conjugated anti-SSEA4 (BioLegend), PE-conjugated anti-KDR (R&D Systems), PECy7-conjugated antiCD31 (BioLegend), PECy7-conjugated anti-CD34 (eBioscience) and FITC-conjugated anti-CD43 (BioLegend).

### Gene expression analysis by RT-semiquantitative PCR

Total RNA was prepared using PureLink RNA micro kits (Invitrogen) in which samples were treated with RNase-free DNase. The reverse transcription of RNA (100 ng-1 μg) into cDNA was performed using random hexamers with Superscript II Reverse Transcriptase (RT) (Life Technologies), according to the manufacturer’s instructions. Real-time quantitative polymerase chain reaction (PCR) was performed on QuantStudio 5 Real-Time PCR Instrument (Applied Biosystems). All experiments were done in triplicate with SYBR-GreenER pPCR SuperMix (Life Technologies), according to the manufacturer’s instructions. Primers (Supplementary File 10) were prepared by Integrated DNA Technologies or Sigma Aldrich. Dilutions of human genomic DNA standards ranging from 100 ng/μl to 10 pg/μl were used to evaluate PCR efficiency of each gene relative to the housekeeping gene *TATA-Box Binding Protein* (*TBP*).

### Microarray analysis

For microarray analysis, 50,000 cells were FACS-sorted directly into Trizol. RNA was extracted from using a miRNeasy Mini Protocol (Qiagen). Samples passing quality control were analyzed using the human Clariom D Assay (ThermoFisher Scientific) and analyzed using Transcriptome Analysis Console (ThermoFisher Scientific) Software and Gene Set Enrichment Analysis (http://software.broadinstitute.org/gsea/index.jsp) software.

### Cell analysis and imaging

For Cytospins, FACS-sorted MKs were spun onto a glass slide and stained with May-Grünwald and Giemsa. Images were obtained on an Olympus BX60 microscope with a 40X objective. An Invitrogen EVOS microscope with a 10x objective was used to image cells in culture.

### Western blots

Cell pellets were resuspended in Laemmli buffer, sonicated for 5 min, and boiled for 5 min at 95 degrees C. Lysates were centrifuged at 10,000 rpm for 5 min at room temperature, and supernatants were used for analysis. Lysate volumes were normalized to cell counts. Samples were run on 4-12% NuPAGE Bis-Tris gels (Invitrogen) and transferred onto nitrocellulose membranes (0.45um pore size, Invitrogen) at 350mA for 90 minutes. Following blocking in 5% milk for 1 h, membranes were incubated with primary antibodies overnight at 4°C. After washing thrice in TBST, membranes were incubated with secondary horseradish peroxidase-conjugate antibodies for 1h at room temperature, washed in TBST thrice, and developed using ECL western blotting substrate (Pierce) and HyBlot CL autoradiography film (Denville Scientific). The following antibodies were used for western blotting: Rabbit anti-TPM1 (D12H4, #3910, Cell Signaling Technologies), Mouse anti-TPM1/TPM2 (15D12.2, MAB2254, Millipore Sigma), Mouse anti-TPM3 (3D5AH3AB4, ab113692, Abcam), (Rabbit anti-TPM4 (AB5449, Millipore Sigma), and Mouse anti-β Actin (A1978, Sigma). Western blot band quantitation was performed using FIJI (Schindelin et al., 2012) (https://fiji.sc/).

### MK activation assay

MKs were pelleted and resuspended in Tyrode’s Salts (Sigma) with 0.1% bovine serum albumin (BSA) containing FITC-conjugated PAC-1 (BD Biosciences), PacBlue-conjugated CD42a (eBioscience) and APC-conjugated CD42b (eBioscience) at a concentration of roughly 100,000 cells per 50μl. Following addition of Convulxin (Enzo Biochem) or Thrombin (Sigma), cells were incubated at room temperature in the dark for 10 min. Cells were then incubated on ice for 10 min. An additional 100μl Tyrode’s Salts containing 0.1% BSA were added and cells were immediately analyzed by flow cytometry.

### Data presentation

Genome-wide SNP Scores were loaded as custom tracks into the UCSC Genome Browser (Kent et al., 2002). Images depicting genomic loci were generated using this tool, as well as Gviz (Hahne and Ivanek, 2016). Other data were created and presented using R, Adobe Illustrator CS6 or GraphPad Prism 6.

### Statistics

Statistical analyses were conducted using R or GraphPad Prism 6.

## Data Availability

All materials, data, code, and associated protocols will be promptly available to readers upon request.

## Acknowledgements

We are grateful for thoughtful suggestions from Drs. Mortimer Poncz, Michele Lambert, Michael Atkinson, Gordon Keller, and members of the Voight laboratory, as well as technical support from Tapan Ganguly and Hetty Rodriguez (University of Pennsylvania Microarray Core Facility), and the Penn Medicine Academic Computing Services. We thank Osheiza Abdulmalik for generous use of his microscope for Cytospin imaging.

This work was supported through R01DK101478 (BFV), a Linda Pechenik Montague Investigator Award (BFV), R01HL130698 (DLF, PG), T32HD043021 (CST), a Children’s Hospital of Philadelphia Neonatal and Perinatal Medicine Fellow’s Research Award (CST), an American Academy of Pediatrics Marshall Klaus Neonatal-Perinatal Research Award (CST) and a Children’s Hospital of Philadelphia Foerderer Award (CST).

## Author Contributions

CST and BFV conceived of this study. CST, CDJ, KL, JAM, DLF, and BFV conducted and/or analyzed experiments. CST and BFV wrote the manuscript. BFV oversaw the work.

## Supplementary Information

### Supplementary Figures and Figure Legends

**Figure 1-figure supplement 1.**
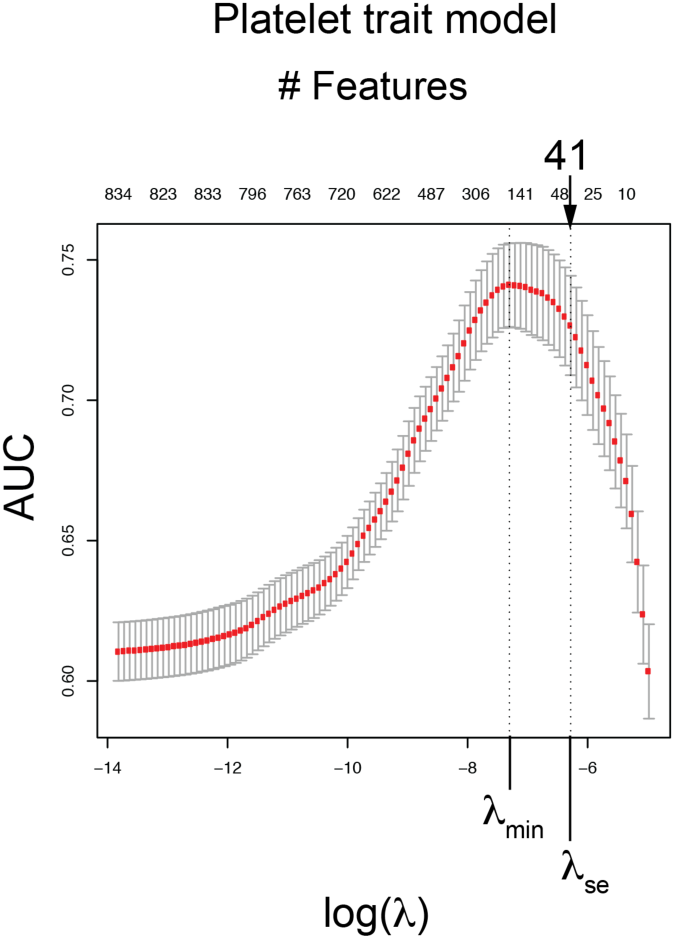
Penalized regression identifies epigenetic features that discriminate platelet trait GWAS SNPs from matched controls. Area under the Receiver Operator Curve (AUC) for platelet trait model. Penalized regression results depicting the regularization parameter (λ) vs. Area under the Receiver Operator Curve (AUC). Top axis shows how many features were identified at each level of λ. Variation in AUC at each λ reflects 10-fold cross-validation. The λ_min_ (model with maximal AUC) and λ_se_ (minimal feature inclusion with AUC within 1 standard error of λ_min_) are shown, with λ_se_ model incorporating the indicated number of features. The final model, with 41 total features, included 38 chromatin features and 3 background characteristics (Distance to Nearest Gene, Minor Allele Frequency, and Number of SNPs in linkage disequilibrium).

**Figure 2-figure supplement 1.**
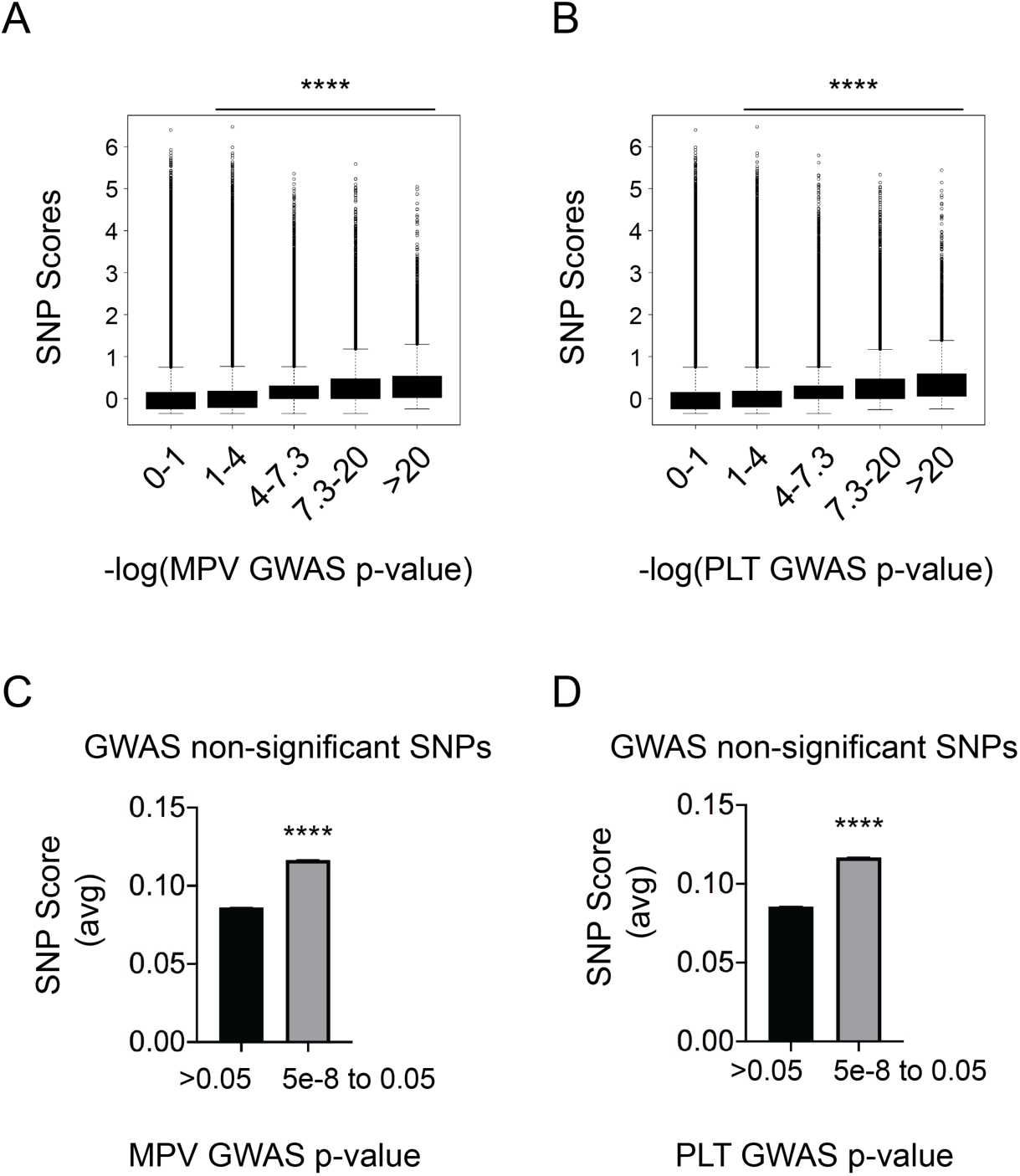
High SNP scores for platelet trait model capture information from sub-genome-wide significant loci. **(A,B)** Higher SNP scores correlate with lower GWAS p-values for variation in (**A**) mean platelet volume (MPV) or (**B**) platelet count (PLT). SNPs were scored genome-wide and Arbitrarily binned –log_10_(p-value) from GWAS MPV or PLT variation was plotted against SNP scores. A value of 7.3 for –log_10_(p-value) correlates with a p-value of 5×10^-8^. Box-and-whisker plots show 25^th^-to-75^th^ percent interval (box) and standard deviation (whiskers). ****p<0.0001 vs Column 1 (ANOVA, Dunnett’s multiple comparison test). Significant linear correlations existed between higher values of –log_10_(p-value) and SNP scores (Pr(>|t|)<2e-16 by linear regression significance test). **(C,D)** SNPs that nearly missed genome-wide significance for (**C**) MPV or (**D**) PLT were enriched for high SNP scores. SNPs that did not meet genome-wide significance were stratified into non-significant (p-value >0.05) and marginally significant (p-value between 5e-8 and 0.05). Bars represent mean±SEM. ****p<0.0001 by Wilcoxon Rank Sum test.

**Figure 2-figure supplement 2.**
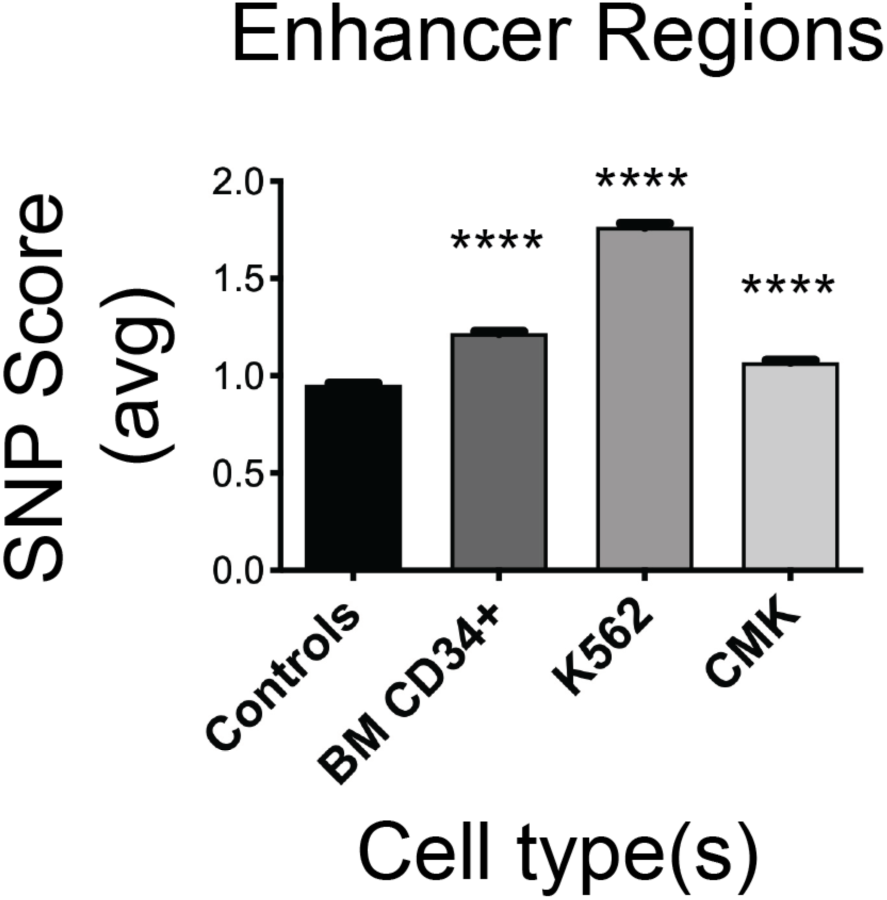
Platelet trait model gives high scores to SNPs marking hematopoietic enhancer regions. Hematopoietic enhancer regions are enriched for high SNP scores based on our platelet trait model. FANTOM5-defined enhancer regions for adult bone marrow (BM) CD34+ (CNhs12553), K562 (human erythroleukemia, CNhs12458), and CMK (human megakaryoblastic leukemia, CNhs11859) hematopoietic cells were compared with enhancer regions from random non-relevant cell types (CNhs11756 from adult pancreas, CNhs14245 from a papillary cell lung adenocarcinoma cell line and CNhs12849 from adult parotid gland). Bars represent mean±SEM. ****p<0.0001 by 1-way ANOVA vs Controls.

**Figure 2-figure supplement 3.**
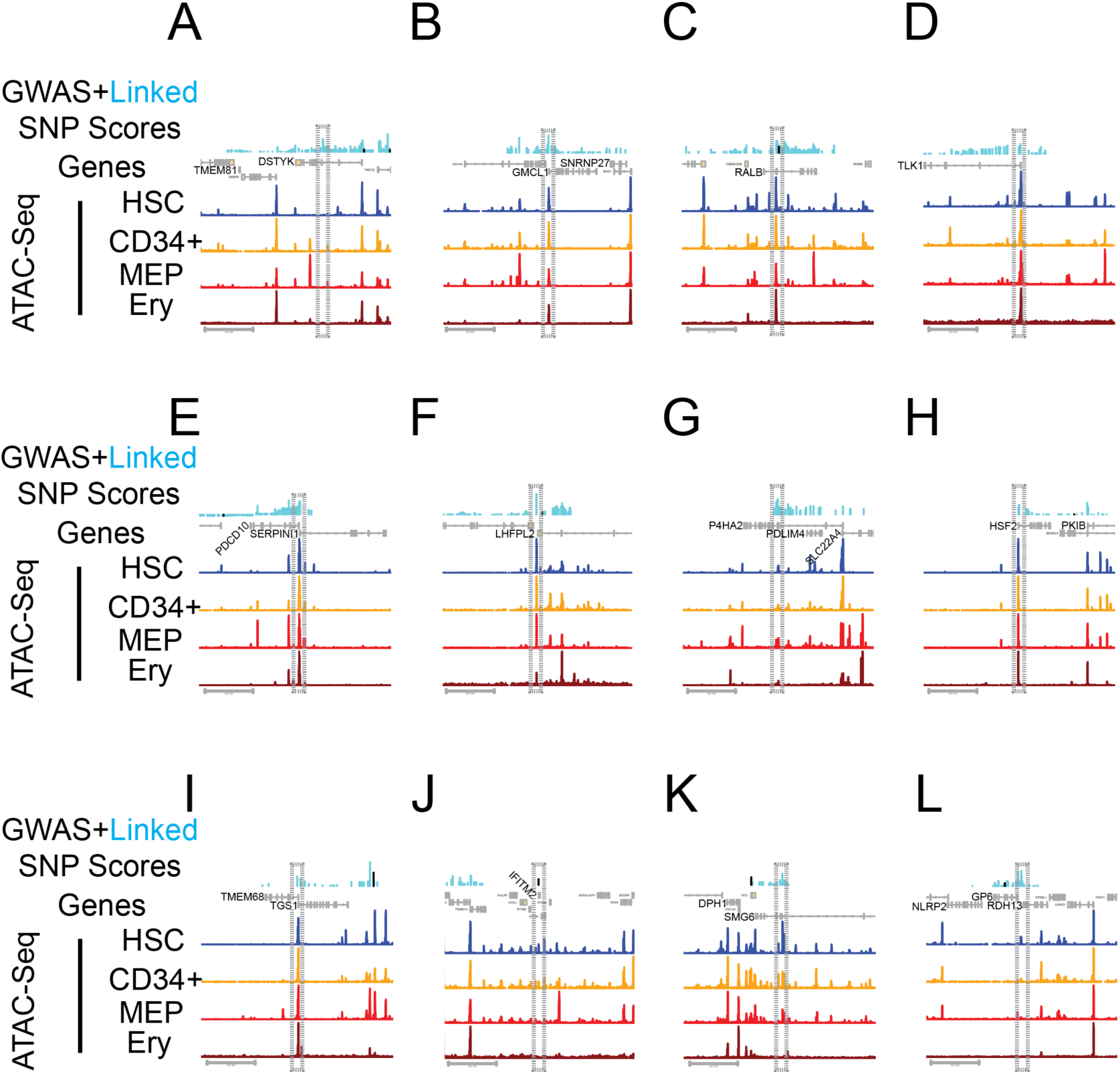
Additional putatively active eQTLs implicated through fine-mapping with LASSO-based SNP scores and by direct overlap with GATA binding sites. In each panel, the top portion shows GWAS SNP in black and linked SNPs (EUR r^2^>0.7) in cyan. Bar heights depict SNP scores. Gene exons are shown in yellow. Accessible chromatin regions (ATAC-Seq) are shown for hematopoietic stem cells (HSC), CD34+ hematopoietic progenitor cells, megakaryocyte-erythroid progenitors (MEP), and erythroblasts (Ery) (Corces et al., 2016). Implicated SNP(s) in region are outlined in gray box, and interesting gene(s) in region are indicated. Note that some SNPs regulate multiple genes, but only nearby regulated genes are boxed and labeled here. **(A)** rs11240368 is an eQTL for *CNTN2* and *TMEM81*. **(B)** rs3771535 is an eQTL for *GMCL1* and *SNRNP27*. **(C)** rs10180681 and rs10180682 are eQTLs for *EPB41L5, PTPN4,* and *RALB*. **(D)** rs9646785 is an eQTL for *GAD1* and *GORASP2*. **(E)** rs6771578 is an eQTL for *PDCD10, SERPINI1,* and *WDR49*. **(F)** rs12652692 is an eQTL for *LHFPL2* and *SCAMP1*. **(G)** rs72793280 is an eQTL for *ACSL6, P4HA2, PDLIM4, SLC22A4,* and *SLC22A5*. **(H)** rs1741820 is an eQTL for *HSF2* and *PKIB*. **(I)** rs13265995 is an eQTL for *LYN, TGS,* and *TMEM68.* **(J)** rs9704108 is an eQTL for *IFITM2.* **(K)** rs2316513 is an eQTL for *DPH1, SMG6,* and *SRR.* **(L)** rs1654439 is an eQTL for *GP6, NLRP2,* and *RDH13.* Scale bars, 50 kb.

**Figure 3-figure supplement 1.**
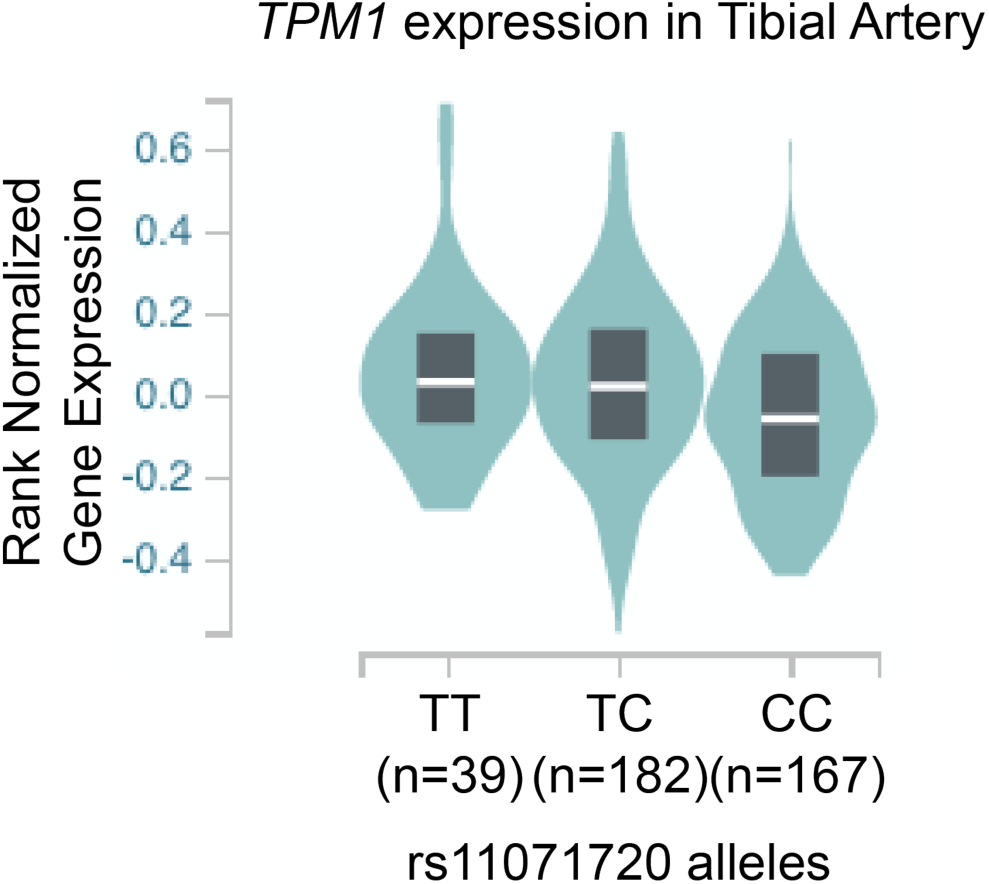
The SNP rs11071720 is an expression quantitative trait locus (eQTL) for *TPM1*. Individuals with the rs11071720 minor ‘C’ allele have decreased *Tropomyosin 1* expression in tibial artery tissue (p= 0.000056, Normalized Enrichment Score= −0.082). Data obtained from GTEx V7 (Ardlie et al., 2015).

**Figure 4-figure supplement 1.**
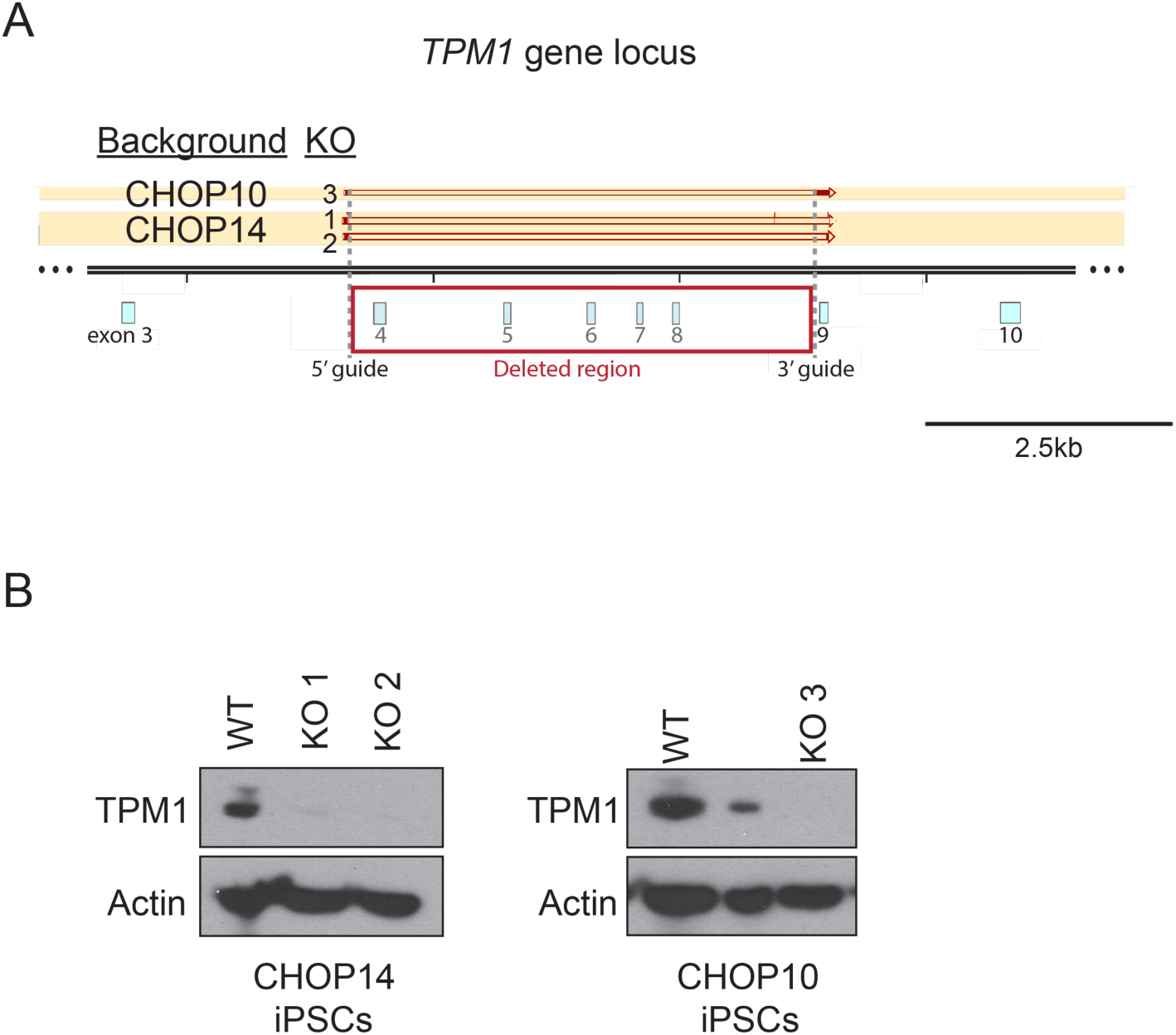
DNA sequencing and western blot confirmation of *TPM1* deletion. **(A)** Shown are *TPM1* exons (numbered light blue boxes) in and around the proposed deletion site. 5’ and 3’ guide RNA sites are marked. Deleted areas in each clone are indicated as ‘empty’ bars, with flanking present DNA in dark red. **(B)** Western blot of CHOP14 or CHOP10 iPSC lysates showing no TP DM1 protein in KO clones. Middle lane in CHOP10 blot depicts a suspected heterozygous clone.

**Figure 4-figure supplement 1.**
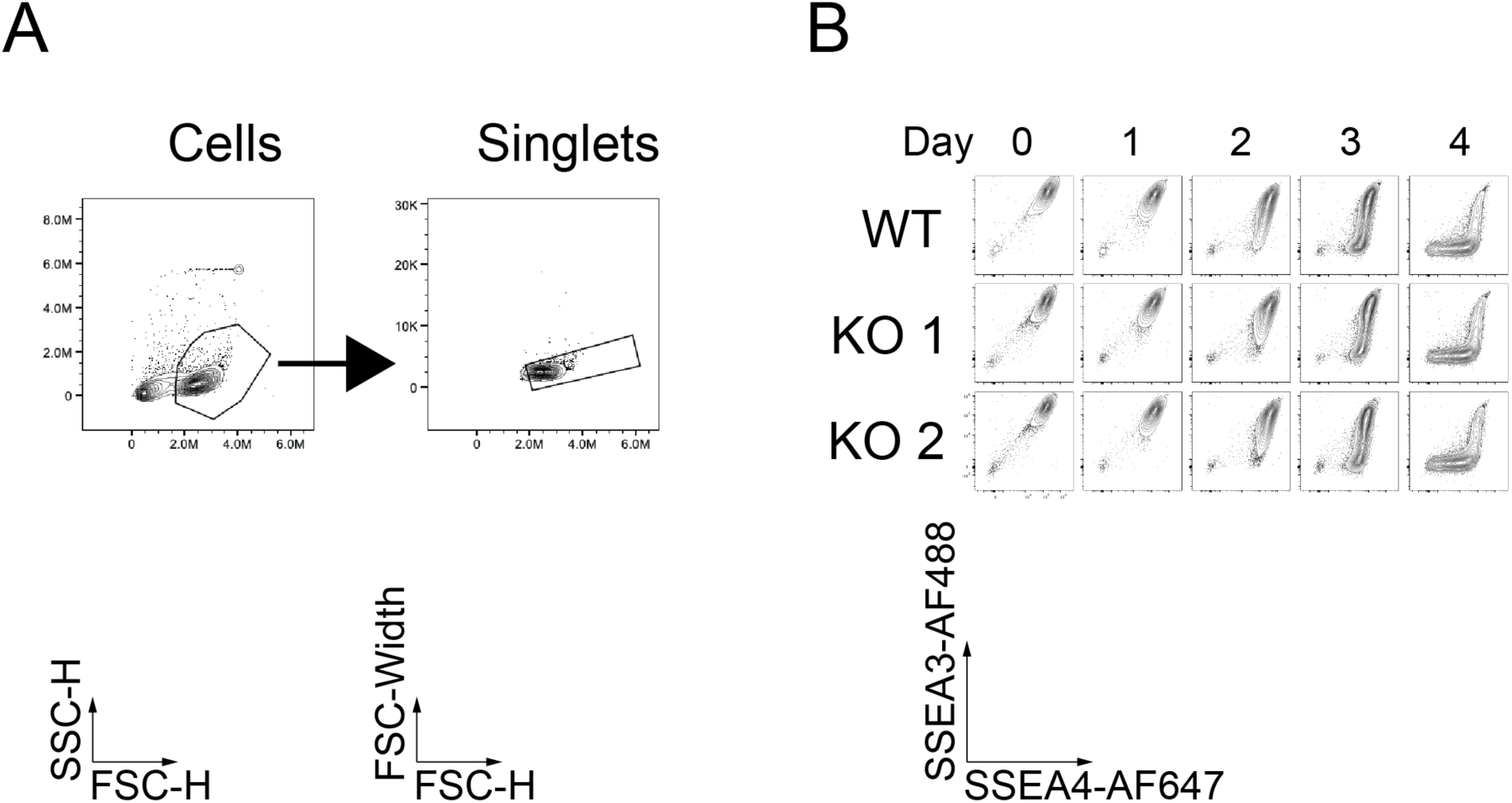
KO cells show normal kinetics of pluripotency marker loss in early differentiation. **(A)** Representative gating strategy for flow cytometry analysis. Singlet cells were analyzed directly for all presented studies. **(B)** On days 0-4, TPM1 KO iPSCs show normal loss of pluripotency markers SSEA3 and SSEA4, with kinetics identical to WT.

**Figure 4-figure supplement 2.**
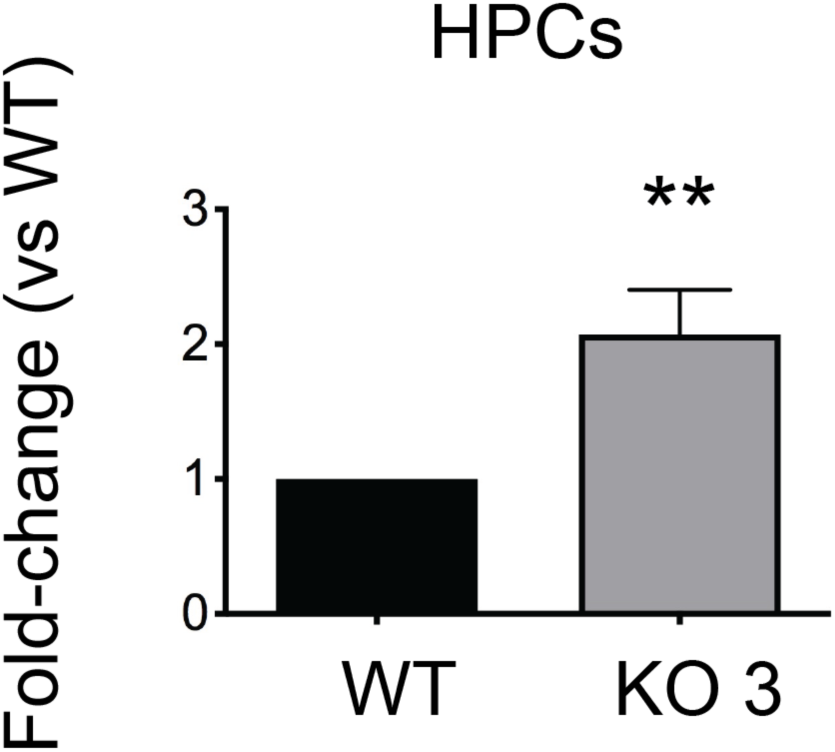
CHOP10-derived *TPM1* KO iPSCs yield more single cells after differentiation. There were more hematopoietic progenitor cells (HPCs, non-adherent single cells) in CHOP10-derived *TPM1* KO clone 3 following 7-8 hematopoietic differentiation. **p<0.01.

**Figure 4-figure supplement 3.**
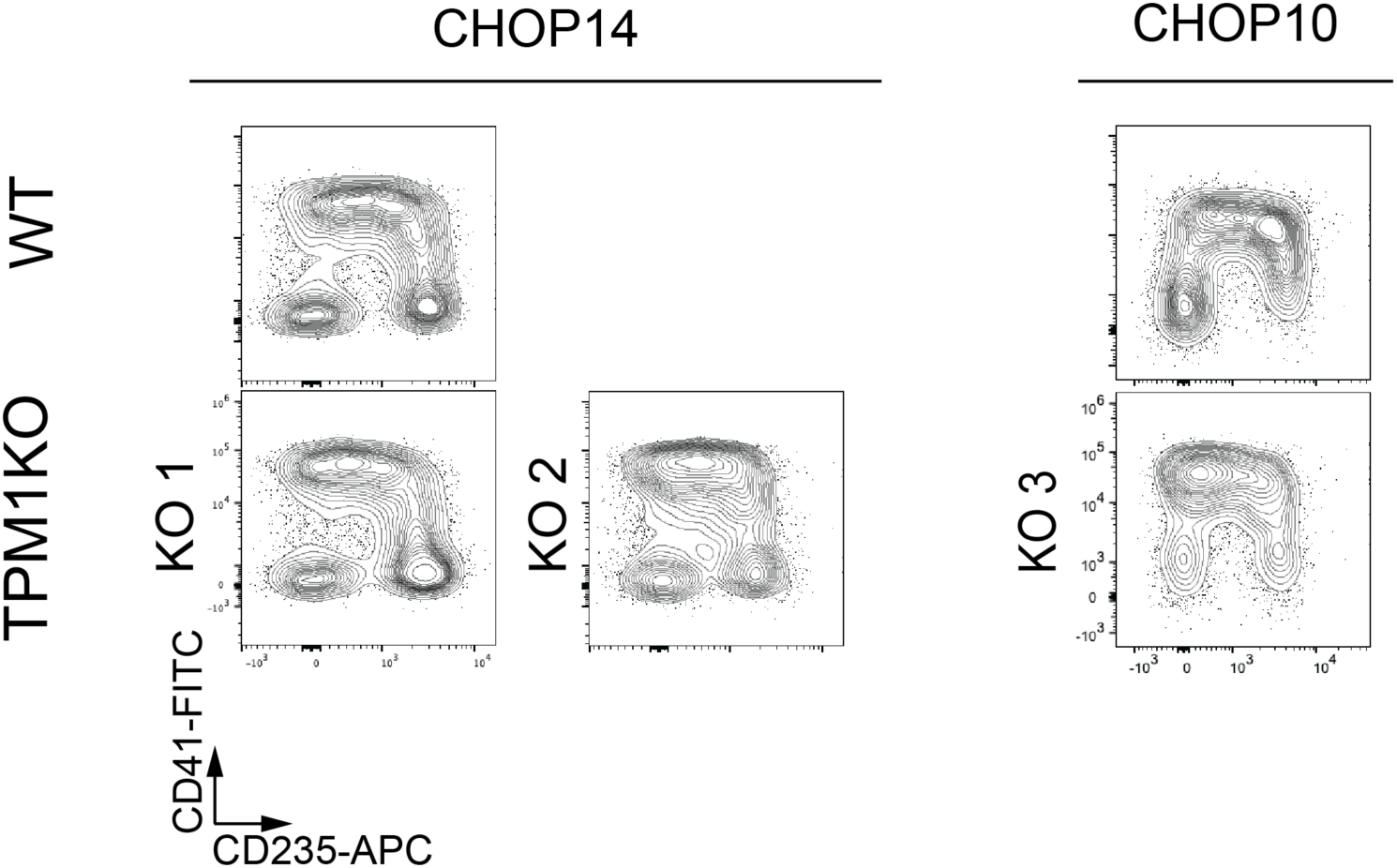
Non-adherent cells (HPCs) from *TPM1* KO cultures show normal cell surface markers. WT and TPM1KO iPSC clones 1-3 all display relatively normal cell surface marker patterns after 9 d differentiation. Multiple experiments show no consistent lineage preference across all clones.

**Figure 5-figure supplement 1.**
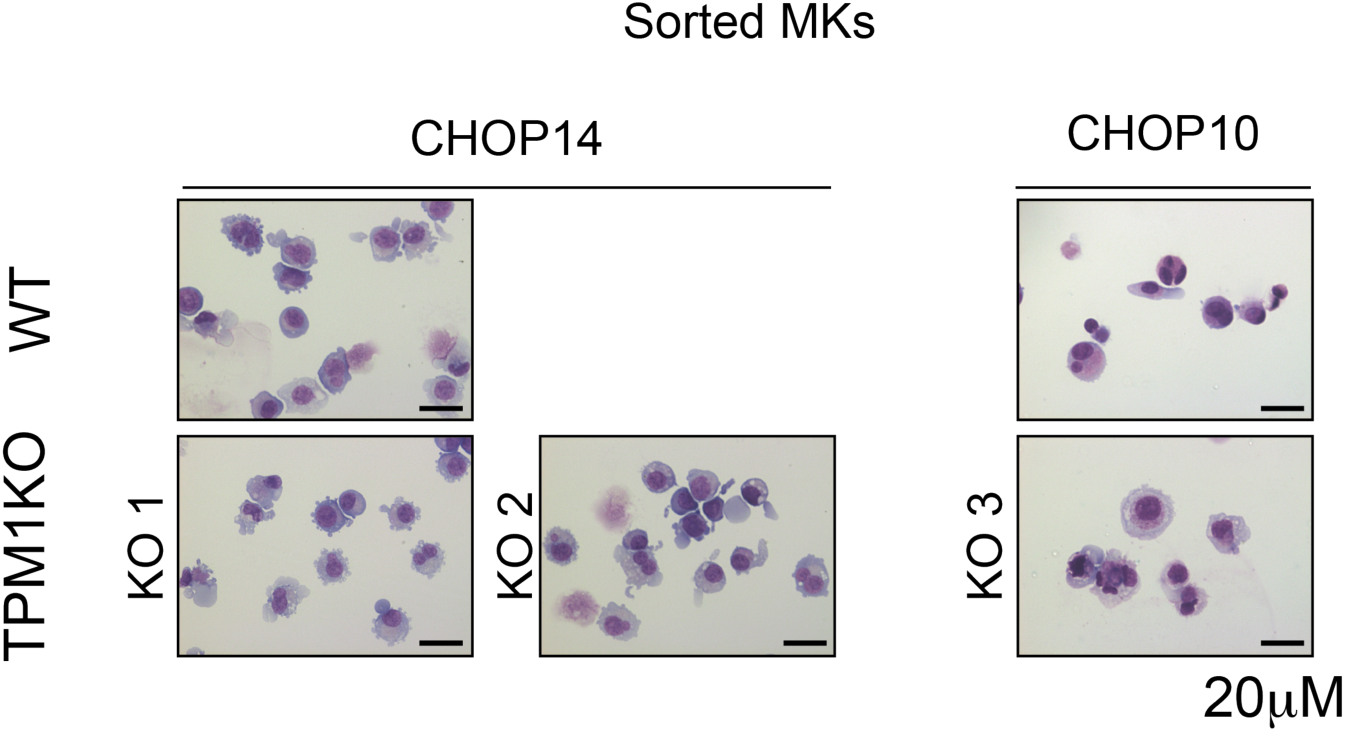
*TPM1* KO MKs have normal morphology. Following 8 d differentiation and 5 d MK expansion culture, wild type (WT) and *TPM1* KO CD41^+^/CD42b^+^ primitive MKs were FACS-sorted and analyzed by Cytospin. Scale bar represents 20μm.

**Figure 5-figure supplement 2.**
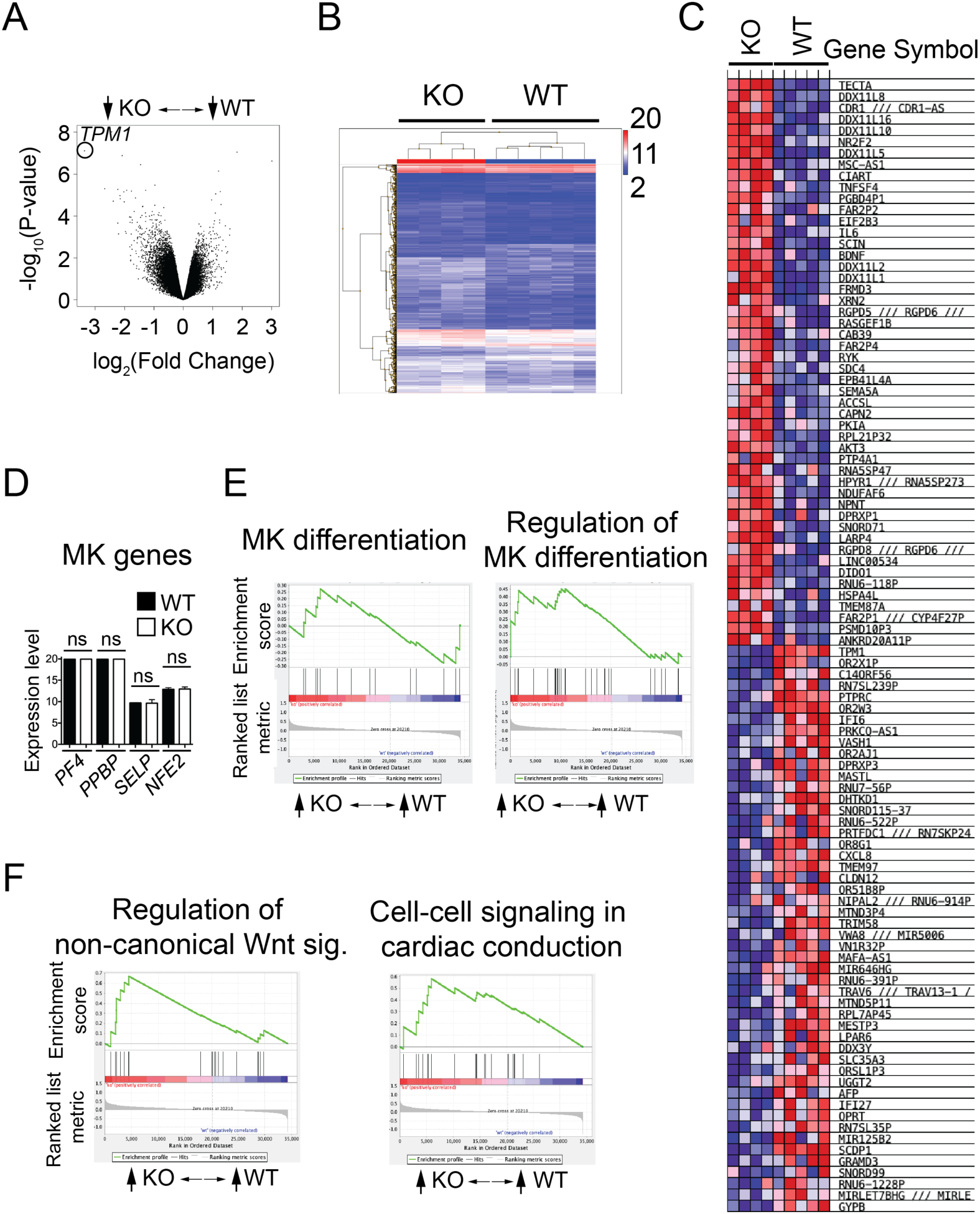
Microarray analysis shows no significant differences in MK genes. **(A)** Volcano plot showing gene expression changes in WT and KO MK microarray analysis. *TPM1* is circled. **(B)** Hierarchical clustering for microarray gene analysis data of FACS-sorted WT and KO MKs. Red, high expression. Blue, low expression. **(C)** Heat map shows the most highly upregulated (top) and downregulated (bottom) genes in KO MKs. **(D)** Expression of representative MK genes are not significantly (ns) changed in WT vs KO MKs. PF4, *Platelet factor 4*. PPBP, *Pro-platelet basic protein*. SELP, *P-selectin*. NFE2, *Nuclear factor erythroid 2*. **(E)** Gene set enrichment analysis (GSEA) for MK pathways were not significantly changed. Shown are GO pathways for MK differentiation (FDR q-value 0.314) and Regulation of MK differentiation (FDR q-value 0.64). **(F)** GSEA plots for select significantly upregulated pathways in KO MKs.

**Figure 5-figure supplement 3.**
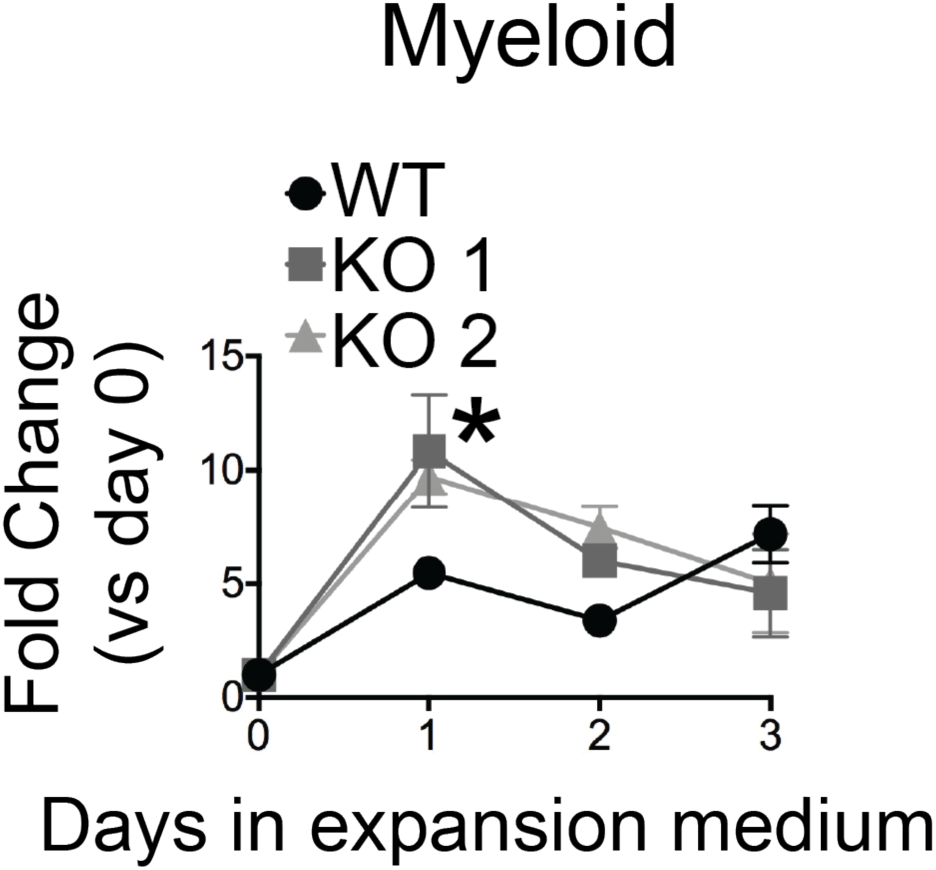
*TPM1KO* HPCs retain normal myeloid lineage expansion potential. HPCs obtained after 8d differentiation were put into lineage expansion media and cultures were analyzed by manual cell counting and flow cytometry over 3-5 d. Mature myeloid cells were CD45^+^. Points represent lineage-specific cell percentage multiplied by total cell count, normalized to cell count on day 0. *p<0.05 by ANOVA vs WT.

**Figure 5-figure supplement 4.**
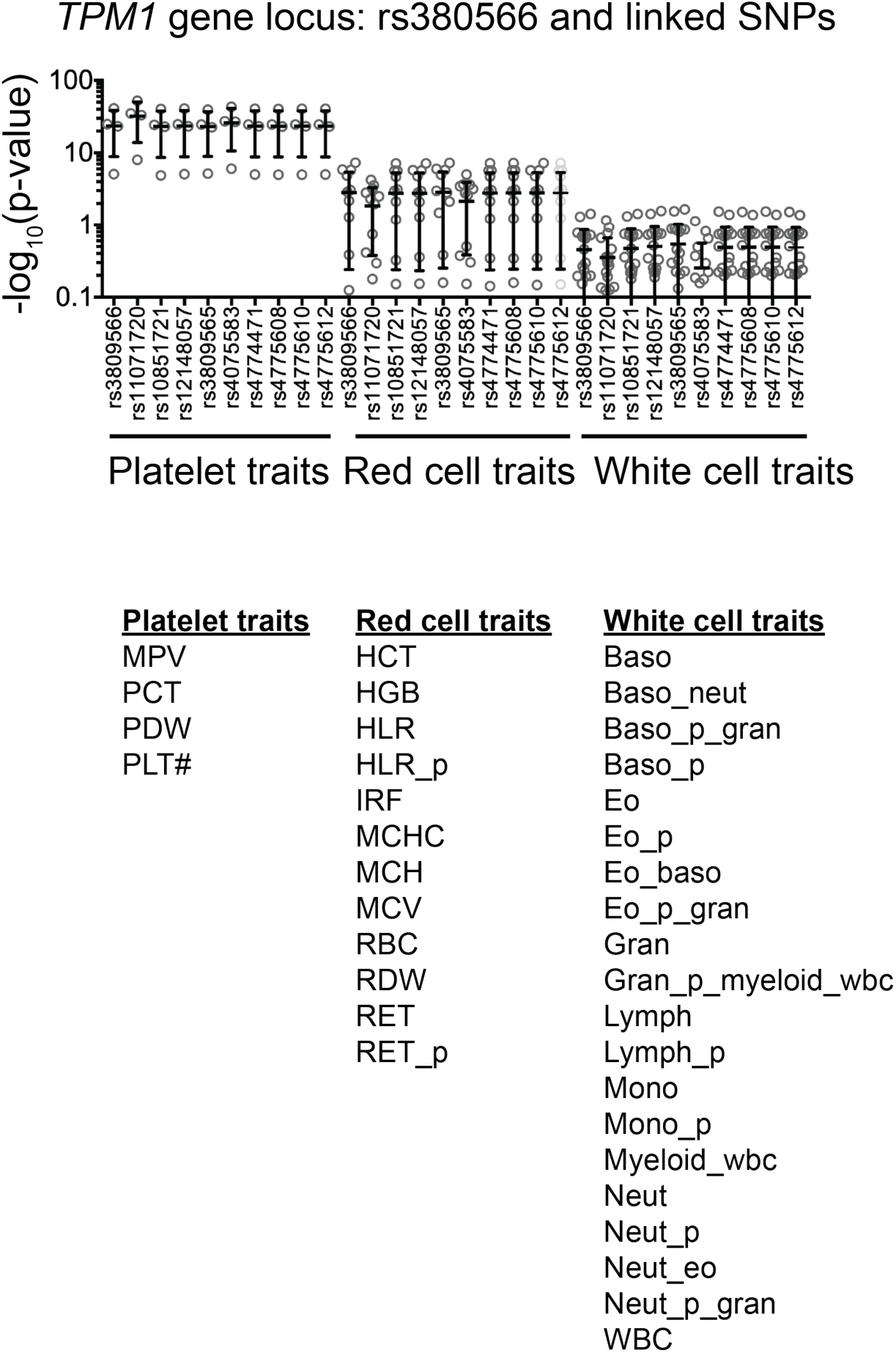
Hematopoietic trait associations of SNPs associated with the *TPM1* gene locus. Aggregated GWAS platelet, red cell, or white cell trait p-values for all SNPs in high LD (EUR r^2^>0.7) with rs380566. The p-values for these SNPs reach genome-wide significance for platelet traits (PLT#, MPV).

**Figure 5-figure supplement 5.**
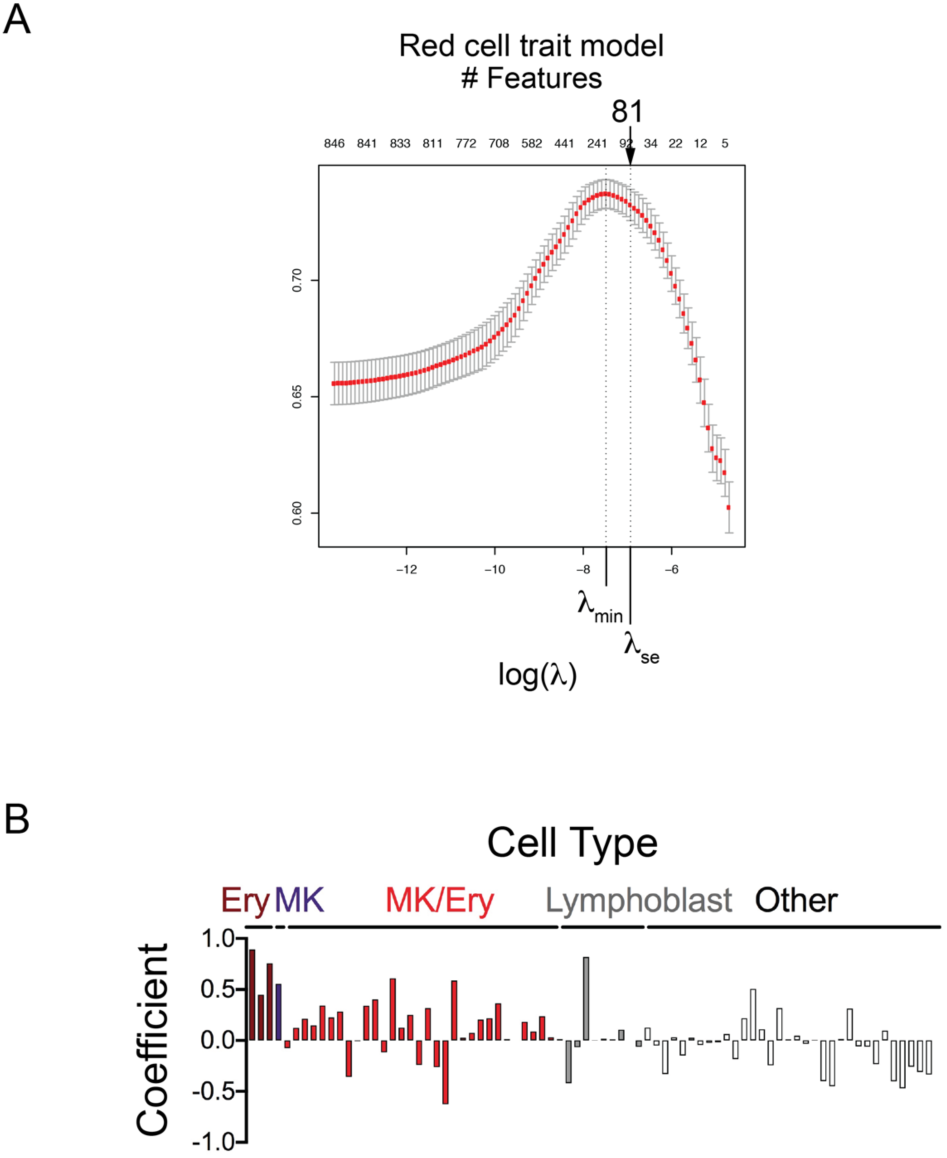
Penalized regression identifies epigenetic features that discriminate red blood cell trait GWAS SNPs from matched controls. **(A)** Area under the Receiver Operator Curve (AUC) for red cell trait model. Penalized regression results depicting the regularization parameter (λ) vs. Area under the Receiver Operator Curve (AUC). Top axis shows how many features were identified at each level of λ. Variation in AUC at each λ reflects 10-fold cross-validation. The λ_min_ (model with maximal AUC) and λ_se_ (minimal feature inclusion with AUC within 1 standard error of λ_min_) are shown, with λ_se_ model incorporating the indicated number of features including background characteristics (Distance to Nearest Gene, Minor Allele Frequency, and Number of SNPs in linkage disequilibrium). **(B)** Penalized regression (LASSO) (Tibshirani, 1996) analysis identified 78 chromatin features from the indicated cell types that best discriminated red cell GWAS SNPs, in addition to 3 background characteristics (distance to nearest gene, minor allele frequency, number of SNPs in linkage disequilibrium). Bar heights are LASSO coefficients, indicating the relative importance of each feature. Ery, peripheral blood derived erythroblasts. MK, primary megakaryocytes. MK/Ery, K562 cells. Lymphoblast, GM12878 or GM12891.

**Figure 5-figure supplement 6.**
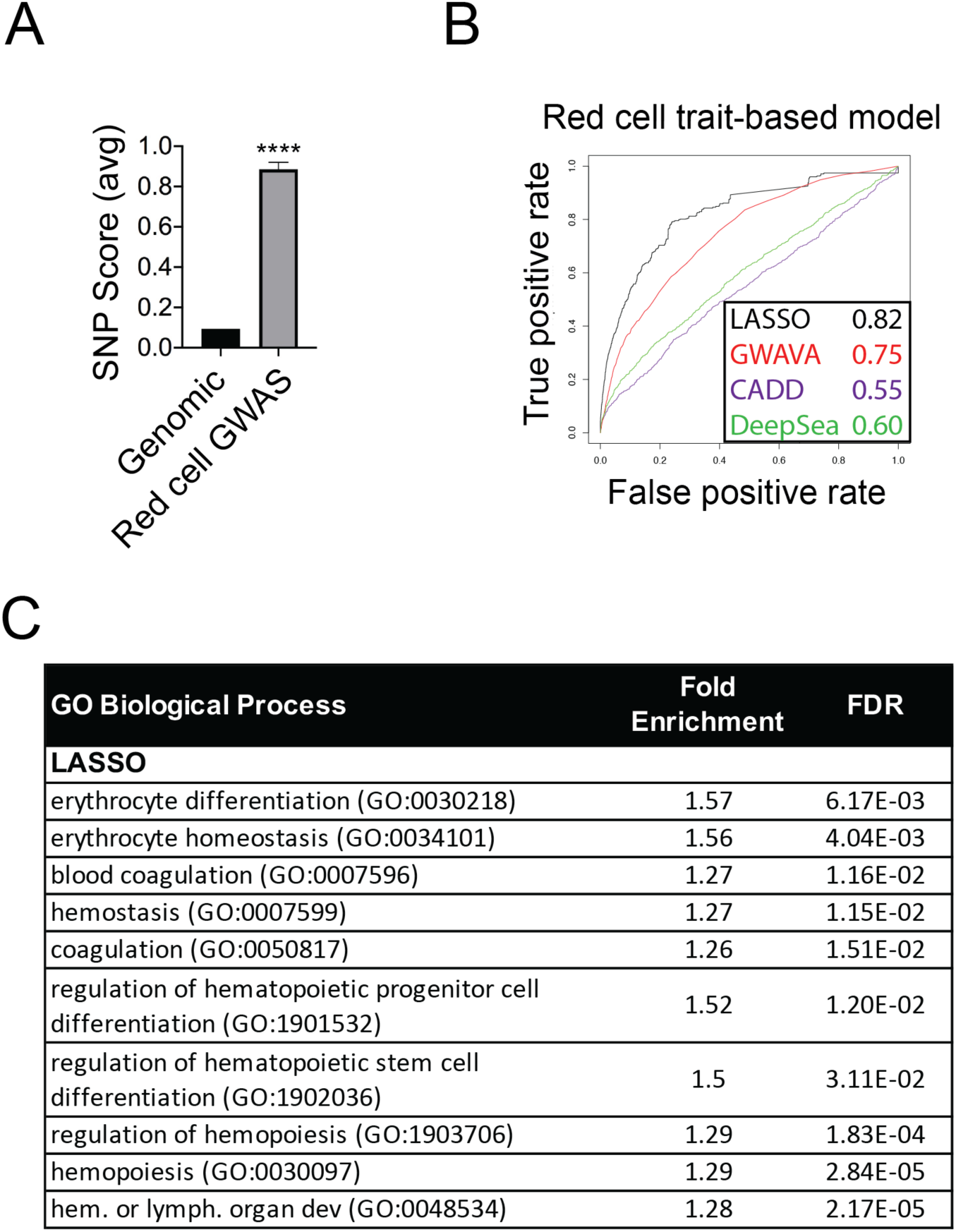
Penalized regression model identifies genes relevant to erythroid and hematopoietic biology. **(A)** SNP scores for red cell trait model training SNPs were significantly higher than genome-wide SNP scores. Bars represent mean+-SEM, ****p<0.0001. **(B)** Performance comparison of our red cell trait model to DeepSEA (Zhou and Troyanskaya, 2015), GWAVA (Ritchie et al., 2014), and CADD (Kircher et al., 2014) for red cell GWAS SNP identification. AUC values are shown in the legend. **(C)** Erythroid and hematopoiesis pathways (Ashburner et al., 2000) identified by the highest-scoring (top 1%) SNPs genome-wide for the red cell model, excluding established red cell trait loci (Astle et al., 2016) (FDR, False Discovery Rate).

**Figure 5-figure supplement 7.**
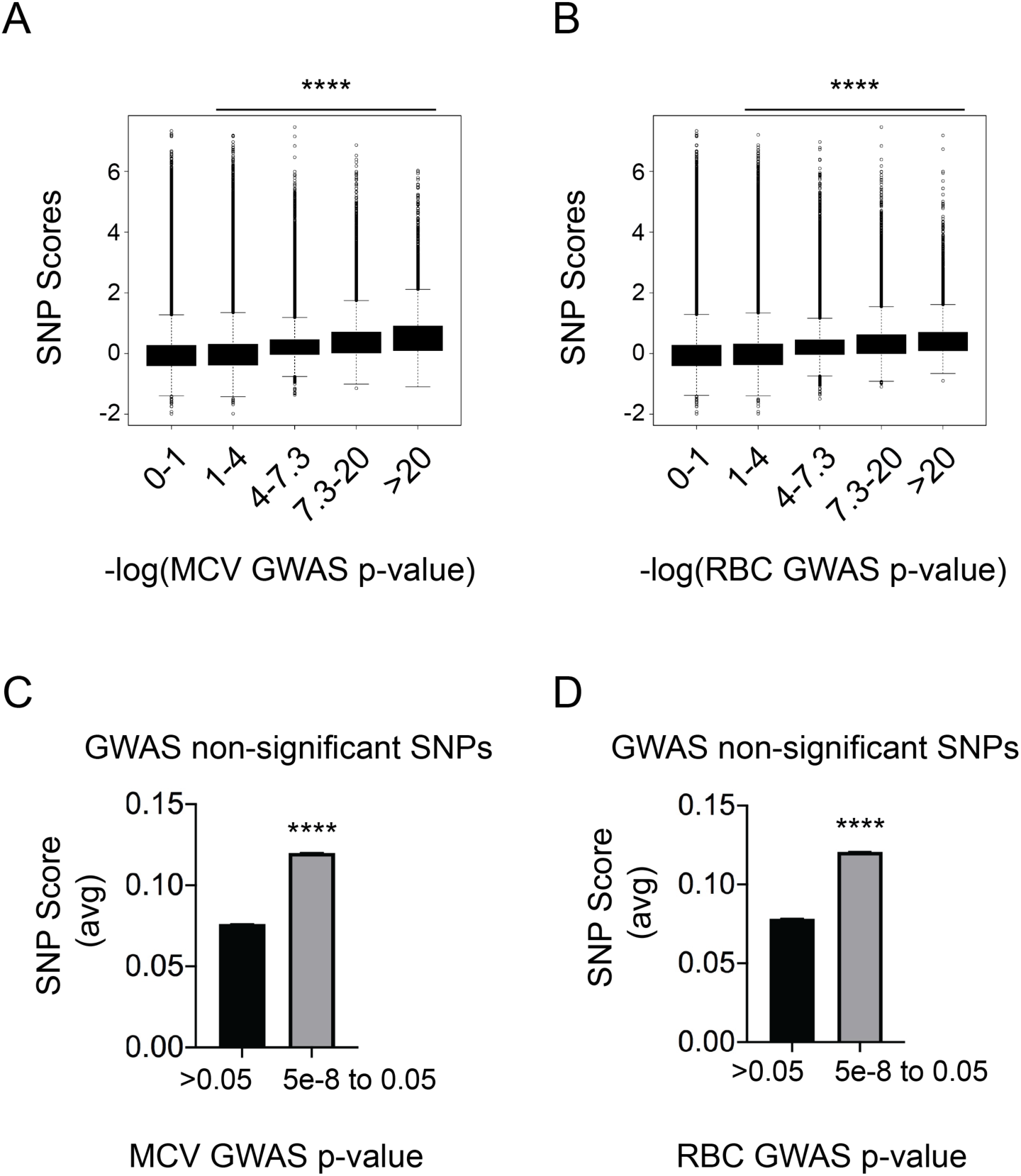
High SNP scores for red cell trait model capture information from sub-genome-wide significant loci. **(A,B)** Higher SNP scores correlate with lower GWAS p-values for variation in (**A**) mean corpuscular volume (MCV) or (**B**) red blood cell count (RBC). SNPs were scored genome-wide and Arbitrarily binned –log_10_(p-value) from GWAS MCV or RBC variation was plotted against SNP scores. A value of 7.3 for –log_10_(p-value) correlates with a p-value of 5×10^-8^. Box-and-whisker plots show 25^th^-to-75^th^ percent interval (red box) and standard deviation (whiskers). ****p<0.0001 vs Column 1 (ANOVA, Dunnett’s multiple comparison test). Significant linear correlations existed between higher values of – log_10_(p-value) and SNP scores (Pr(>|t|)<2e-16 by linear regression significance test). **(C,D)** SNPs missed genome-wide significance for (**C**) MCV or (**D**) RBC were enriched for high SNP scores. SNPs that did not meet genome-wide significance were stratified into non-significant (p-value >0.05) and marginally significant (p-value between 5e-8 and 0.05). Bars represent mean±SEM. ****p<0.0001 by Wilcoxon Rank Sum test.

## Supplementary File Legends

**Supplementary File 1. The 860 chromatin feature tracks included in our LASSO analysis**. These data were obtained from ENCODE (Feingold et al., 2004), ChromHMM (Ernst and Kellis, 2012), and two publications describing analyses of primary human MK cells (Paul et al., 2013; Tijssen et al., 2011).

**Supplementary File 2. Chromatin features and coefficients comprising our penalized regression-based platelet scoring model.**

**Supplementary File 3. Gene Ontology pathways that were significantly enriched in the top 1% of SNPs, as defined by platelet model scores.** Presented pathways had false discovery rate (FDR) < 5%.

**Supplementary File 4. Gene Ontology pathways that were significantly enriched in the top 1% of SNPs, as defined by GWAVA scores.** Presented pathways had false discovery rate (FDR) < 5%.

**Supplementary File 5. Gene Ontology pathways that were significantly enriched in the top 1% of SNPs, as defined by CADD scores.** Presented pathways had false discovery rate (FDR) < 5%.

**Supplementary File 6. Dysregulated molecular pathways in *TPM1KO* MKs**. FACS-sorted MKs were analyzed by microarray, and gene set enrichment was performed. Upregulated Gene Ontology (Ashburner et al., 2000) pathways with FDR<25% are shown. There were no significantly downregulated pathways. *GO, Gene Ontology. NES, nominal enrichment score. FDR, false discovery rate.

**Supplementary File 7. Chromatin features and coefficients comprising our penalized regression-based red cell scoring model.**

**Supplementary File 8. Gene Ontology pathways that were significantly enriched in the top 1% of SNPs, as defined by red cell model scores.** Presented pathways had false discovery rate (FDR) < 5%.

**Supplementary File 9. Penalized regression-based fine-mapping identifies eQTLs in established platelet and/or red cell trait GWAS loci that overlie GATA binding sites.** Listed SNPs are within platelet or red cell trait GWAS LD blocks (EUR r^2^>0.7), scored in the top 5% by *both* our platelet trait and red cell models, overlap canonical or near-canonical GATA binding sites, and are eQTLs for at least 1 gene (GTEx V7) (Ardlie et al., 2015). Associated GWAVA (Ritchie et al., 2014) scores are present, if available. SNP rsIDs and locations refer to hg19 genome.

**Supplementary File 10.** Semi-quantitative RT-PCR primers used in this study.

